# Breast tumor Insulin-like growth factor receptor regulates cell adhesion and metastasis: Alignment of mouse single cell and human breast cancer transcriptomics

**DOI:** 10.1101/2021.08.31.458283

**Authors:** Alison E. Obr, Yung-Jun Chang, Virginia Ciliento, Alexander Lemenze, Krystopher Maingrette, Joseph J. Bulatowicz, Quan Shang, Emily Gallagher, Derek LeRoith, Teresa L. Wood

**Affiliations:** Department of Pharmacology, Physiology & Neuroscience, New Jersey Medical School, Rutgers University, Newark, New Jersey; Office of Advance Research Computing, Rutgers University, Piscataway, New Jersey; Department of Pathology, New Jersey Medical School, Rutgers University, Newark, New Jersey; Division of Endocrinology, Diabetes and Bone Diseases, The Samuel Bronfman Department of Medicine, Icahn Sinai School of Medicine at Mt. Sinai, New York, New York 10029

**Keywords:** IGF-1R, breast cancer, cell adhesion, metastasis, cadherin, METABRIC

## Abstract

The acquisition of a metastatic phenotype is the critical event that determines patient survival from breast cancer. Several receptor tyrosine kinases have functions both in promoting and inhibiting metastasis in breast tumors. Although the insulin-like growth factor 1 receptor (IGF-1R) has been considered a target for inhibition in breast cancer, low levels of IGF-1R expression are associated with worse overall patient survival. To determine how reduced IGF-1R impacts tumor phenotype, we used weighted gene correlation network analysis (WGCNA) of METABRIC patient data and identified gene modules specific to cell cycle, adhesion, and immune cell signaling inversely correlated with IGF-1R expression in human breast cancers. Integration of human patient data with data from mouse tumors revealed similar pathways necessary for promoting metastasis in basal-like tumors with reduced signaling or expression of the IGF-1R. Functional analyses revealed the basis for the enhanced metastatic phenotype including alterations in E- and P-cadherins.

## Main

Metastasis is the leading cause of cancer patient death. Several individual genes and associated cellular pathways contribute to a metastatic phenotype but the mechanisms that cause some tumors to become metastatic are still poorly understood. Receptor tyrosine kinases (RTKs) have been implicated in promoting metastatic properties in tumor cells. RTK domain mutations are not a prominent feature in most cancers; instead, RTK expression level is the general driver of tumorigenesis and metastasis (1–4). A well-known RTK that has a prominent role in a subclass of breast cancers and has been the focus for successful cancer therapeutics is HER2. However, targeting several other RTKs including the epidermal growth factor receptor (EGFR) and the insulin-like growth factor receptor (IGF-1R) in breast tumors has been mostly unsuccessful (2, 5, 6). The emerging theme for these receptors is their context-dependent functions that change whether they are growth-promoting or growth-inhibiting in the primary tumor or metastatic environment.

Expression of IGF-1R has been implicated in tumor oncogenesis by promoting tumor cell proliferation and survival (7–9). Due to this oncogenic function, several IGF- 1R inhibitors have been developed and used in clinical trials. While IGF-1R was a clear target, the inhibitors were largely unsuccessful in the clinic (5, 6). There is now clear evidence that the IGF-1R also has tumor or metastasis suppressive functions; IGF-1R expression in breast tumors correlates with positive overall patient survival and a more differentiated tumor phenotype (10–12). Consistent with these data, recent reports using the TCGA and METABRIC patient databases have revealed low IGF-1R expression is associated with undifferentiated, triple-negative breast cancer (TNBC) and worse overall survival (13, 14).

In the present study, we utilized the METABRIC patient database and single-cell RNA sequencing of two IGF-1R loss-of-function mouse tumor models to uncover how IGF-1R signaling regulates intrinsic epithelial cell signaling to suppress metastasis. We identify key pathways necessary for promoting metastasis including upregulation of immune cell activation signals, cell cycle dysregulation, and altered cell adherence and show that IGF-1R is required to maintain a metastasis suppressive tumor microenvironment. We further show that adherence between luminal and basal tumor cells is necessary for tumor growth at the secondary site and that reduced IGF-1R signaling in tumor epithelial cells dysregulates E- and P-cadherin resulting in reduced cell adhesion.

## Methods

### Animal Models

All animal protocols were approved by the Rutgers University Institutional Animal Care and Use Committee (Newark, NJ) and all experiments were managed in accordance with the NIH guidelines for the care and use of laboratory animals. Animal care was provided by the veterinary staff of the division of animal resources in the New Jersey Medical School Cancer Center of Rutgers Biomedical Health Sciences. The *MMTV-Wnt1* line on an FVB background [FVB.Cg-Tg(Wnt1)1Hev/J] was obtained as a gift from Dr. Yi Li. The *MMTV-Wnt1//MMTV-dnIgf1r* (referred to here as DN-Wnt1) line was described previously (15).

Mice carrying floxed alleles of exon 3 of the *Igf1r* gene (16) were bred with a keratin 8 (K8)-Cre^ERT^ transgenic line (JAX stock #017947) (17) and with the *MMTV- Wnt1* transgenic line to produce female mice that were homozygous for the *Igf1r* floxed alleles and hemizygous for both the K8-Cre^ERT^ and *MMTV-Wnt1* transgenes referred to as K8iKOR-Wnt1 mice (see Supplemental Methods).

### Mammary Tumor Epithelial Cell Dissociation

Tumor mammary epithelial cells (MECs) were isolated from Wnt1, DN-Wnt1, and K8iKOR-Wnt1 mice similarly to our prior study (15). Whole tumors were excised and dissociated with the gentleMACs tissue dissociator (130-093-235, protocol m_TDK2) and mouse specific tumor dissociation kit (Miltenyi, 130-096-730). Organoids that retain basement membrane attachments were trypsinized (0.05% Trypsin-EDTA, Gibco) and filtered with a 40 μm cell strainer (BD Biosciences) to isolate a single cell suspension of dissociated tumor MECs (see also Supp. Methods). Isolated tumor MECs were counted with a hemocytometer for flow cytometry, FACS, tail vein injections (TVIs), *in vitro* adhesion assays, and cell culture assays.

### Tail vein injection (TVI) of primary tumor epithelial cells

Tumor MECs were isolated as described above (n=4) and injected at 1×10^6^ cells/200ul PBS (unsorted) or 0.25×10^6^ cells/200ul (sorted) into the tail vein of 6-week-old eGFP mice (n=4). Animals were perfused with 3% PFA and lungs were harvested at 1-, 3-, 6-, 8-, and 12-weeks post injections. After perfusion, harvested lungs were drop-fixed in 4% PFA overnight and dehydrated with 70% EtOH for paraffin embedding. For TVIs with flow sorted tumor MECs (n=3), 0.25×10^6^ luminal (CD24^+^/CD29^lo^), basal (CD24^+^/CD29^hi^), or unsorted epithelial cells/200ul PBS were injected into eGFP mice (n=3); lungs were harvested and processed 1-week post injection (wpi).

### RNA isolation and real-time quantitative PCR

RNA was purified from whole tumor and sorted tumor epithelial cells according to the manufacturer’s protocol (Qiagen). RNA concentration and quality was assayed with the NanoDrop ND-1000 (Thermo Scientific). Sorted tumor epithelial cell cDNA was transcribed according to manufacturer’s protocol using SuperScript II (Invitrogen) from total RNA (200 ng). Samples were run in technical triplicate to determine relative gene expression by real-time quantitative PCR (qRT-PCR) detected with SsoAdvanced Universal SYBR Green Supermix (BioRad) using the BioRad CFX96 real-time PCR machine according to manufacturer’s instructions. Transcript levels were normalized to glyceraldehyde-3-phosphate dehydrogenase (GAPDH) or Gusb for mouse and ß-actin for human, and data were analyzed using the Q-Gene software (BioTechniques Software Library) (18). For detection of the *Igf1r*-deleted allele in isolated cell populations, we amplified the corresponding fragment from the coding region of the messenger RNA specific for the inactivated receptor allele. The forward primer annealed with exon 2 and the reverse primer spanned the knockout-specific splice junction between exons 2 and 4. Primer oligonucleotide pairs for qRT-PCR are provided (Supp. Table 1).

### Histology and Immunofluorescence

Tumor tissues and lungs from animals with primary tumors (n=4 per genotype) were drop-fixed in 4% paraformaldehyde (PFA), embedded in paraffin, and sectioned at 7 µm. Lung sections from animals with primary tumors or from TVIs were used for hematoxylin and eosin staining. Tumor and lung sections were processed for antigen retrieval for immunofluorescence (IF) as described previously (19). Tissue sections were immunostained with primary antibodies: E-cadherin (1:100; Invitrogen, ECCD-2), cytokeratin-8 (1:100; TROMA-I, DSHB), cytokeratin-14 (1:250; Invitrogen, PA5-16722), phospho-Histone H3 (Ser10) (1:200; Cell Signaling, D2C8 XP), P-cadherin (1:100; Invitrogen, MA1-2003), and Ki67 (1:100; Vector Labs, VP-K451) and with species-specific fluorochrome-conjugated secondary antibodies (1:500, Invitrogen).

A Keyence BZ-X all-in-one fluorescence microscope with BZ- scientific imaging processing software (Keyence) was used to capture images. At least 5 individual fields were captured at 20X or 40X magnification from tumor sections (n=4 per genotype). For thicker sections, a z-stack range was acquired and the focus analysis was utilized to obtain the deconvoluted image.

### Tumor epithelial cell *in vitro* adhesion assays

Primary tumors were dissociated as described above and incubated in tissue culture on collagen coated plates for 10 hours. Culture media (DMEM/F12, 5% FBS, insulin (5 μg/mL), EGF (5 ng/mL), hydrocortisone (1 μg/mL), 0.1% gentimicin) was removed and cells in suspension were fixed on slides using a cytospin (Shandon Cytospin 3) for 10 minutes at 1500 rpm for immunofluorescence (IF). Cells attached to the collagen matrix were fixed with 4% PFA for 10 minutes at room temperature for IF analysis or lysed with RLT buffer (Qiagen) for RNA isolation and qRT-PCR analysis as described above.

For IF, cells were processed for staining as previously described (20). Cells were stained with primary antibodies: cytokeratin-8 (1:100; TROMA-I, DSHB) and cytokeratin- 14 (1:250; Invitrogen, PA5-16722) and with species-specific fluorochrome-conjugated secondary antibodies (1:500, Invitrogen). To visualize cell nuclei, cells were stained with DAPI (1:10,000 in PBS). Images were captured as described above and cells were manually counted using ImageJ.

### Single-cell RNA sequencing

Whole Wnt1 (tamoxifen injected, Cre negative), DN-Wnt1, and K8iKOR-Wnt1 tumors were dissociated as described above except tumor cells were filtered with a 70 μm filter directly after dissociation to collect single cells from the entire tumor. Cells were captured using the 10X Chromium system (10X Genomics) and sequenced with the NextSeq 500 (Illumina). Analysis is described in Supp. Methods.

### WGCNA analysis of METABRIC data for gene module identification

The data generated from 1981 patients within the METABRIC project (21) was used in this investigation. These data were accessed through Synapse (synapse.sagebase.org), including normalized expression data and clinical feature measurements. The associated expression Z scores were downloaded from cBioPortal (https://www.cbioportal.org/). The method of weighted gene co-expression network analysis (WGCNA) was used to identify gene modules with significant statistical association to the phenotypic trait including patient age, tumor size, tumor grade, cancer subtype, and IGF-1R expression. The description of clinical feature coding and gene correlation analysis is found in Supp. Methods.

### Ingenuity Pathway Analysis (IPA)

#### scRNA-seq

Differentially expressed gene sets were identified from the DN-Wnt1 and K8iKOR-Wnt1 compared to Wnt1 mouse tumors for each whole tumor and epithelial cell specific cluster determined from scRNA-seq as described above. These differentially expressed genes were used for IPA pathway enrichment and graphical summary analysis. The top 5 pathways based on significance were plotted by percent genes altered in each pathway. Graphical summaries were generated using the top pathways, cell functions, and target genes identified from differentially expressed genes (DN-Wnt1 vs. Wnt1; K8iKOR-Wnt1 vs. Wnt1) in each cluster.

#### WGCNA METABRIC analysis

Gene names and expression levels identified from highly correlative co-expression gene modules as described in Supp. Methods were uploaded into the IPA software (Qiagen) and analyzed for pathway enrichment. The top 5 pathways based on log-fold change significance for each module were plotted by percent of total genes up- and down-regulated in each pathway.

#### Comparison Analysis

Whole tumor gene changes were compared to ME genes where the output is pathway alterations. Here, exact genes were not completely similar, but pathways were comparable.

### Statistics

All graphical data were expressed as the mean ± SEM. Statistical comparisons were carried out by GraphPad Prism9 software. The Student’s *t*-test or non-parametric Mann-Whitney U test was used for two-group comparisons. Specific comparisons are described in figure legends when necessary. For multiple variable analysis, the One-Way ANOVA with Tukey’s Multiple Comparison post-hoc test was performed. For the tumor growth curve and *in vitro* adhesion analysis, the non-linear regression least squares regression for slope best fit was used to compare differences between each line. The Chi-Square test was used to determine differences between genotypes in the metastasis table. Power calculations were performed based on pilot data to determine the number of tumor samples necessary using a 2-sided hypothesis test, an *α* = 0.0025, and 80% power.

## Results

### Low levels of IGF-1R correlate with a metastatic gene signature in breast cancer

Recent analysis of TCGA and METABRIC databases have revealed IGF-1R expression is reduced in TNBC (13, 14). Furthermore, low levels of IGF-1R predict worse overall patient survival across all breast cancer subtypes (14, 22). Our previous studies reported IGF-1R expression levels in human tumors are inversely correlated with several key target genes that alter the tumor microenvironment (14) . These prior expression analyses of human breast tumors with low IGF-1R were performed on genes we identified as dysregulated with reduced IGF-1R signaling and associated with increased metastasis in our mouse tumor model (14, 23). The findings from human and mouse support the hypothesis that low expression of IGF-1R could be used to identify gene signatures associated with aggressive breast cancers. To independently stratify genes correlated with either low or high IGF-1R expression in human breast cancers, we performed a global unbiased weighted gene co-expression network analysis (WGCNA) utilizing the METABRIC database to identify gene expression modules associated with IGF-1R expression Z-score, referred to as IGF1R gene set 1 (IGF1R- GS1; Supp. Fig. 2).

Due to the large number of genes in the IGF1R-GS1, we refined our WGCNA analyses to limit the original data set to those genes with the strongest positive or negative correlation to IGF-1R expression (Fig. 1a). In this refined gene set (IGF1R- GS2), we identified four gene co-expression modules significantly correlated with low IGF-1R (correlation score ≤ -0.25), all of which were also associated with high tumor grade and three of which were associated with TNBC. One module significantly associated with high IGF-1R (correlation 0.61) was also associated with ER+/PR+ breast cancers and low tumor grade (Fig. 1a).

**Figure 1.**
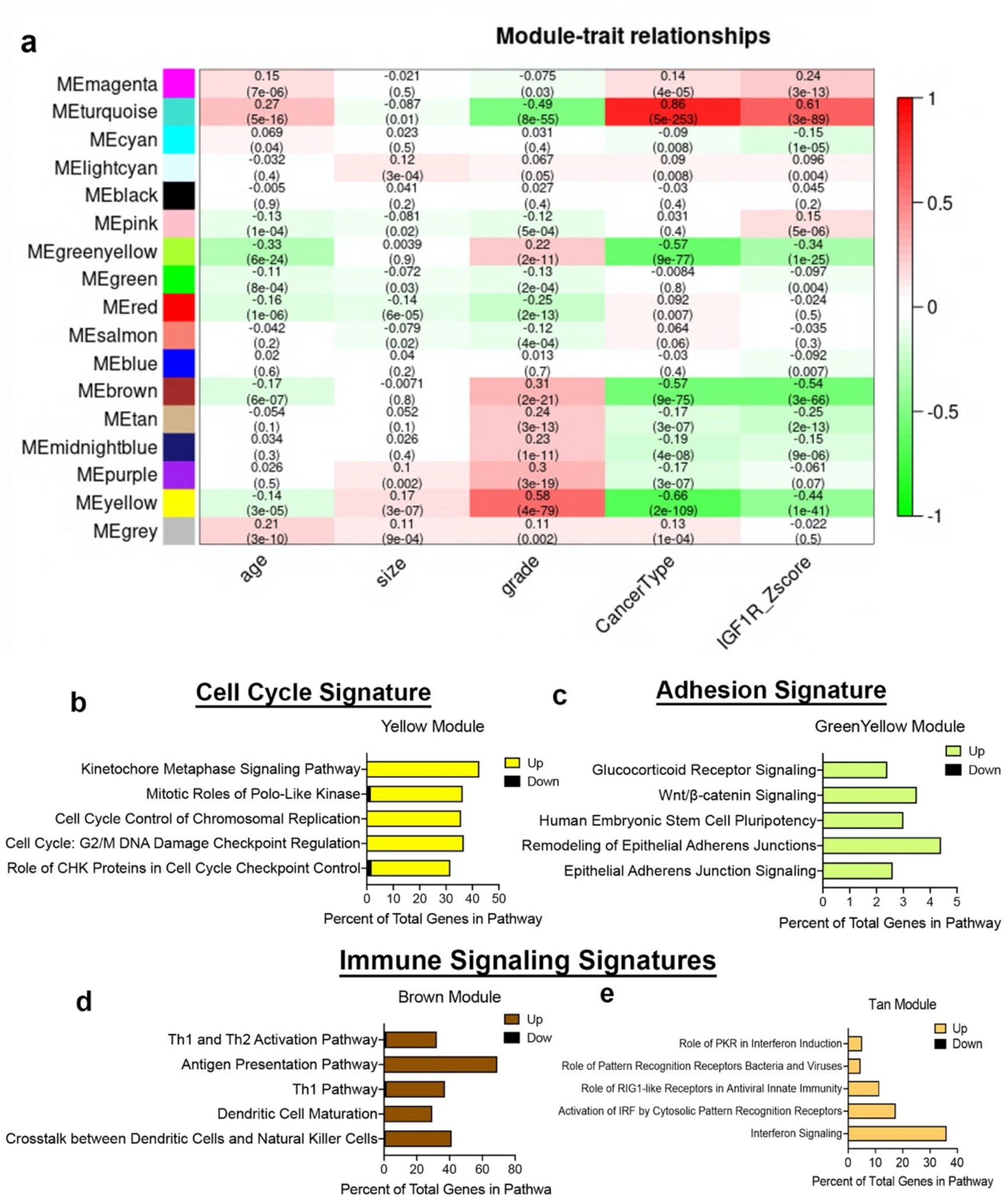
Defining gene signatures associated with IGF-1R expression and tumor phenotype in human BCs. **a.** Table of refined integrated WGCNA (IGF1R-GS2) showing module and clinical trait association. Each row corresponds to a module eigengene (ME), each column to a clinical measurement. Each cell contains the corresponding correlation and p-value (in parentheses). The table is color-coded by correlation according to the color legend. Green < 0 for negative correlation; Red > 0, for positive correlation. **b-e.** Top 5 pathways identified by ingenuity pathway analysis (IPA) revealing key signatures in 4 modules inversely correlated with IGF-1R expression. (yellow module=cell cycle signature, greenyellow module=adhesion signature, brown and tan modules=immune signaling signatures).

Ingenuity pathway analysis (IPA) of IGF1R-GS2 for the pathways associated with the lowest IGF-1R Z-scores revealed genes involved in control of cell cycle checkpoint regulation and chromosome replication, (yellow, Cell Cycle Signature; Fig. 1b) and in epithelial adherens junctions (green-yellow, Adhesion Signature; Fig. 1c). The two additional modules associated with low IGF-1R contained genes involved in immune cell signaling (brown, tan; Fig. 1d,e). Taken together, we hypothesize reduced IGF-1R in breast tumors alters both intrinsic tumor epithelial cell pathways and extrinsic immune microenvironment signatures to promote metastasis.

A major question that arises from the METABRIC WGCNA is whether there is a causative relationship between IGF-1R expression and associated gene alterations and, ultimately, phenotype of breast cancer. We published previously that low IGF-1R expression predicts poor patient survival across all breast cancer subtypes (14, 23) suggesting negative functional consequences from loss of IGF-1R expression. Our goal in this study was to use mouse models to test the hypothesis from the human data that low IGF-1R in breast tumors directly contributes to a metastatic phenotype through dysregulated expression of specific cellular pathways.

### Mammary epithelial cell specific IGF-1R deletion promotes Wnt1 driven tumor metastasis

To test how loss of IGF-1R alters the primary tumor phenotype, we made use of two distinct mouse models. In one model developed previously in our lab, IGF-1R function is reduced through expression of a dominant-negative human *Igf1r* transgene (*dnIgf1r*) in the *MMTV-Wnt1* (Wnt1) basal-like breast cancer tumor model (DN-Wnt1; (23)). In this mouse line, the loss of IGF-1R function results in decreased tumor latency and increased lung metastases, while tumor growth is unchanged (23). To model human breast cancers with low IGF-1R expression, we also generated a mammary luminal epithelial lineage-specific *Igf1r* knockout mouse driven from a tamoxifen-inducible Keratin 8 (K8) promoter, referred to as the K8iKOR line (Fig. 2a; see Supp. Methods). Loss of *Igf1r* was verified in mammary epithelial cells (MECs) isolated from hyperplastic glands in 16-week-old virgin K8iKOR-Wnt1 mice compared to control, Wnt1 mice (Supp. Fig. 2a). Decreased *Igf1r* gene expression was maintained in tumors of the K8iKOR-Wnt1 line (Supp. Fig. 2b).

**Figure 2.**
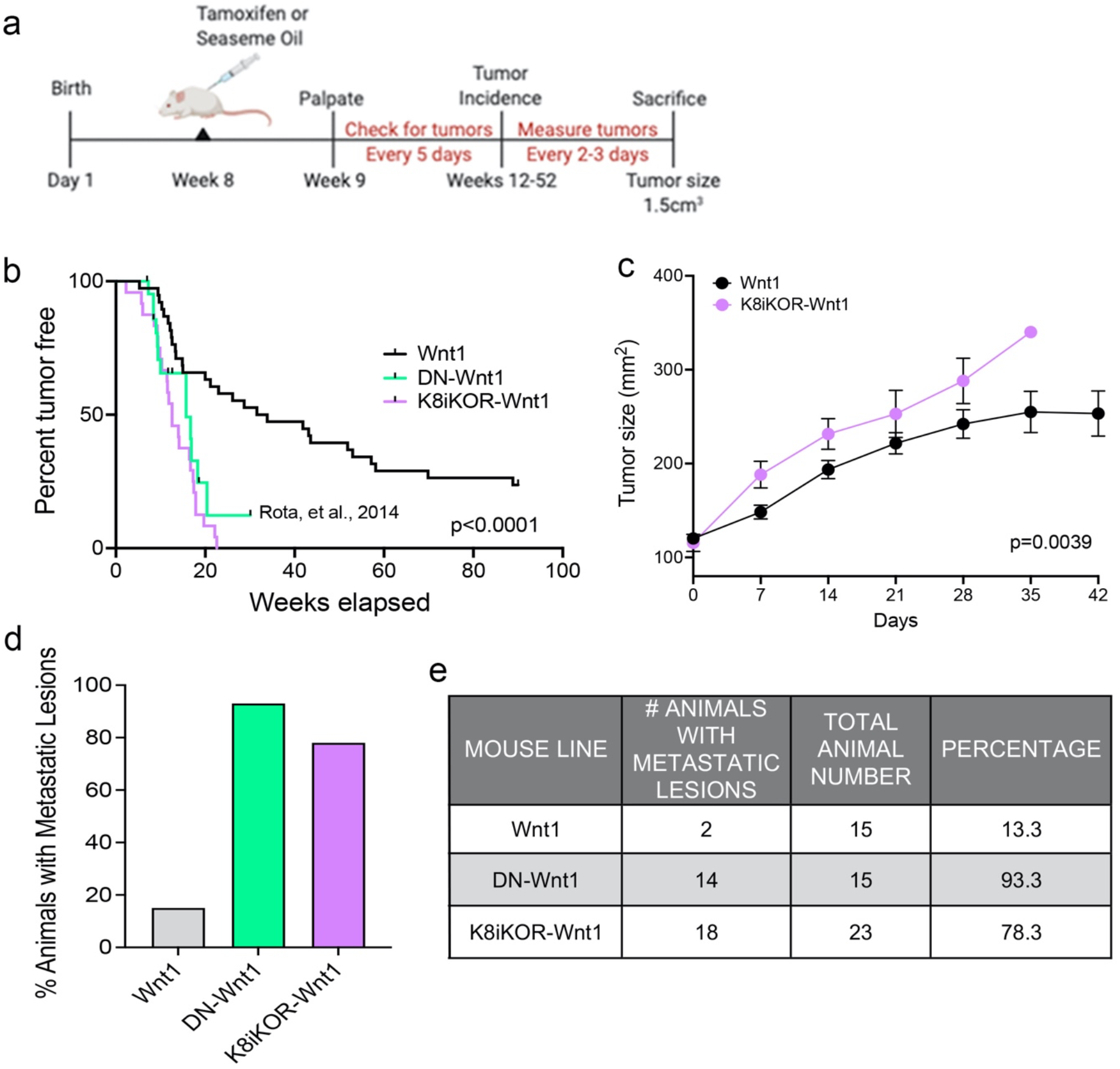
Luminal loss of IGF-1R decreases tumor latency and increases metastasis. **a.** Schematic for luminal lineage IGF-1R knockout. **b.** Latency curve for tumor development in Wnt1, DN-Wnt1, and K8iKOR-Wnt1 animals. For K8iKOR-Wnt1 animals, tumor latency is weeks post tamoxifen injection. *Statistic:* Mann-Whitney test **c.** Growth curve after tumors arise until time of euthanization. *Statistic:* Non-linear regression best fit for line slopes **d-e.** Graph of the percentage of animals **(d)** and table of number of animals **(e)** with metastatic lesions after establishment of a primary tumor. *Table Statistic*: Chi-square test; p=0.0251 for Wnt1 vs. DN-Wnt1 and K8iKOR-Wnt1. For Wnt1 controls, vehicle and tamoxifen injected animals were combined as the phenotypes were equivalent.

To determine the effects of luminal epithelial specific *Igf1r* gene deletion in Wnt1- driven mammary tumorigenesis, we assessed tumor latency rates in the K8iKOR-Wnt1 mouse line compared to the control Wnt1 line and to our prior tumor latency data on the DN-Wnt1 mouse line (23). The mean tumor latency of Wnt1 mice was consistent with previous reports (24, 25), where 50% of control Wnt1 animals formed palpable tumors at 41.7 weeks of age (Fig. 2b). Tumor latency was significantly decreased in K8iKOR- Wnt1 mice (12.5 weeks after tamoxifen injection, p<0.0001) (Fig. 2b) similar to the DN- Wnt1 mouse line as previously reported (16.6 weeks, p<0.0001) (Fig. 2b) (23). Once tumors formed, tumor growth was significantly increased in K8iKOR-Wnt1 compared to control Wnt1 tumors (Fig. 2c). These data indicate that decreased expression of *Igf1r* in luminal epithelial cells accelerates tumor initiation as well as tumor growth in the context of elevated Wnt signaling.

Although the Wnt1 tumors model a basal-like TNBC, these tumors have low metastatic potential (24). In contrast, loss of luminal epithelial *Igf1r* in the Wnt1 tumors significantly increased the percentage of animals with lung micrometastases similar to the metastatic rate in the DN-Wnt1 mice (Fig. 2d,e). Thus, either reduced *Igf1r* expression or reduced IGF-1R function in mammary epithelium promotes metastasis of the primary Wnt1 tumor cells.

### Single-cell sequencing of mammary tumors to analyze epithelial IGF-1R function in regulating tumor cell heterogeneity

Reduced IGF-1R by function or by expression results in increased tumor metastasis in the mouse models and aligns with human survival data indicating an inverse relationship between IGF-1R expression and overall patient survival (14). The mechanisms by which IGF-1R regulates tumor metastasis could include intrinsic epithelial mesenchymal transition (EMT) changes as well as alterations to the tumor microenvironment (TME) secondary to the genetic changes in the tumor epithelium. To reveal underlying mechanisms and cell population changes downstream of alterations in IGF-1R, we performed single cell RNA-sequencing (scRNA-seq) on the DN-Wnt1, K8iKOR-Wnt1 and Wnt1 tumors. We initially analyzed scRNA-seq of the whole tumor to profile changes in tumor cell populations when IGF-1R is either reduced or attenuated in the tumor epithelium. Wnt1 control, DN-Wnt1 and K8iKOR-Wnt1 tumor cells were plotted together resulting in 16 separate tumor cell populations (Fig. 3a). These populations were further defined using cell specific markers resulting in the following distinct cell populations: 6 epithelial, 2 fibroblast (CAF), 6 macrophage/monocyte (MAC), 1 T-cell, and 1 endothelial (Fig. 3b,c, Supp. Fig. 3). Overall, loss of IGF-1R expression or function resulted in decreased macrophage and T cell populations and increased CAF populations (Fig. 3d). Ingenuity pathway analysis (IPA) supports the conclusion that loss of IGF-1R function promotes an immune evasive TME (Fig. 3d,e, Supp. Fig. 4). For example, while the cell number is unchanged in MAC Cluster 0 from DN-Wnt1 and K8iKOR-Wnt1 tumors compared to Wnt1, the immune function pathways are altered with downregulation of genes involved in immune cell activation, antigen presentation, cell adhesion, and infiltration (Fig. 3e, Supp. Fig. 4a).

**Figure 3.**
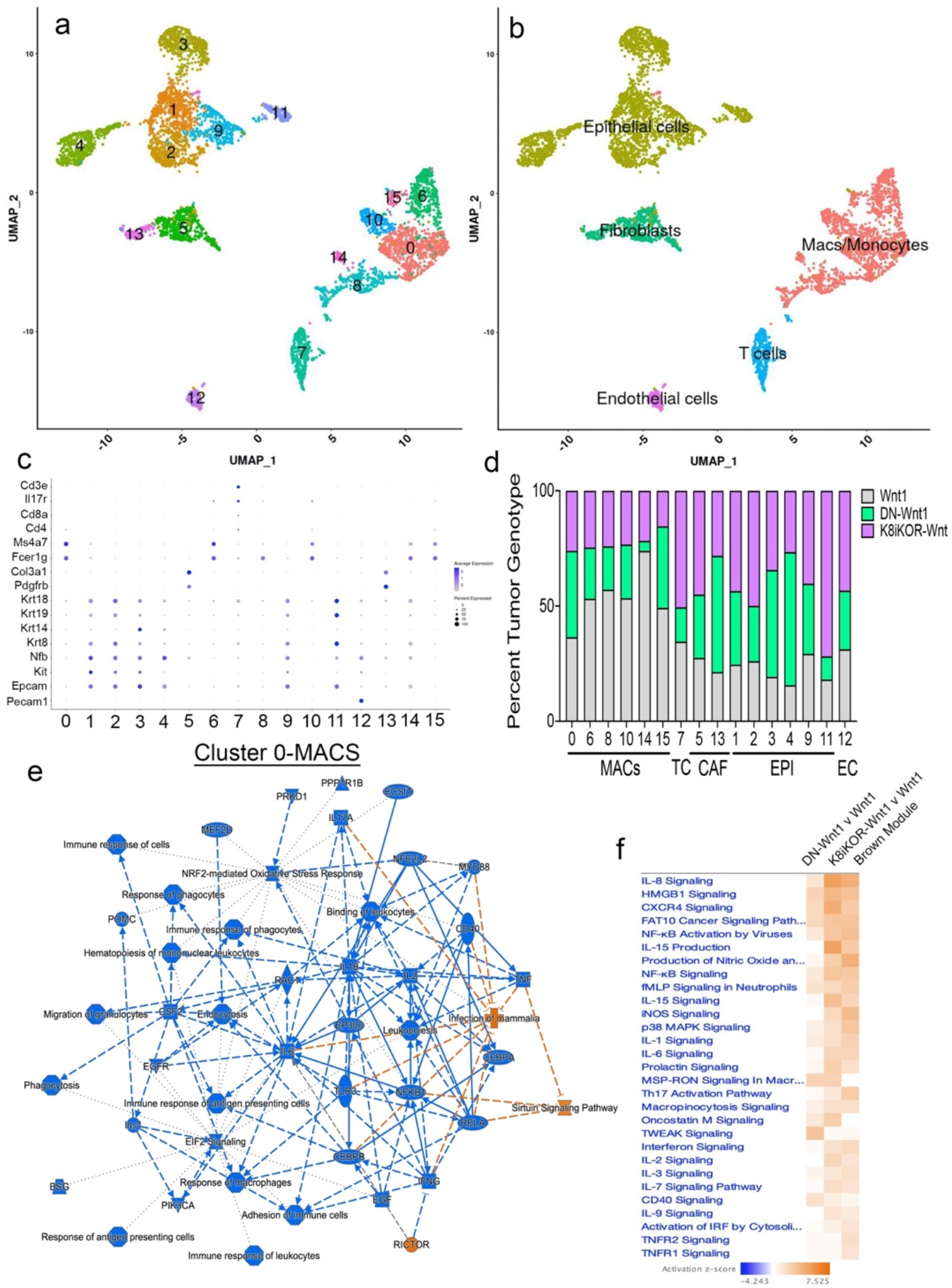
Identifying mammary tumor heterogeneity by single cell RNA-sequencing. **a.** Uniform Manifold Approximation and Projection (UMAP) plot of cells from Wnt1, DN-Wnt1, and K8iKOR-Wnt1 tumors resulting in 16 individual clusters. **b.** UMAP plot with identification of cluster cell types defined by known markers. **c.** Dot plot of cell markers. **d.** Percent tumor genotype graph for each cluster. Clusters are ordered by identified tumor cells. MAC and T-cell populations were generally decreased in DN- Wnt1 and K8iKOR-Wnt1 tumors. CAF populations were expanded in DN-Wnt1 and K8iKOR-Wnt1 tumors. (MACS=monocytes/macrophages, TC=T cells, CAF=fibroblasts, EPI=epithelial cells, EC=endothelial cells) **e.** IPA graphical summary of top pathway alterations in DN-Wnt1 compared to Wnt1 tumors from Cluster 0 (MACs). Blue=downregulated; orange=upregulated. **f.** IPA canonical pathways heat map of DN- Wnt1 and K8iKOR-Wnt1 compared to Wnt1 tumors and the METABRIC brown (immune signaling signature) module.

Alignment of the immune signature module from the METABRIC data analysis (Fig. 1d) revealed several immune signaling pathways similarly associated with human patient tumors with low IGF-1R expression and mouse tumors with reduced IGF-1R function or expression (Fig. 3f). Interestingly, the pathways upregulated in both patient and mouse tumors with reduced IGF-1R are important for response to stress signaling and immune cell evasion supporting our prior findings that loss of IGF-1R promotes cell stress in human breast cancer cells (14).

### Reduced IGF-1R alters the microenvironment to promote metastasis

Our findings from the single cell analyses of the tumor modules suggest two possible non-exclusive hypotheses: increased metastasis with decreased IGF-1R is due to 1) microenvironment alterations, and/or 2) epithelial cell intrinsic alterations. To directly test intrinsic tumor cell invasive capacity *in vivo* we performed tail vein injections (TVI) with isolated tumor epithelial cells. We injected Wnt1, DN-Wnt1 or K8iKOR-Wnt1 tumor epithelial cells into tail veins of eGFP mice and analyzed lungs for metastases 1- week post injection (1 wpi; Fig. 4a,b). Tumor cells from all three genotypes formed the same number of micrometastases 1 wpi suggesting reduced IGF-1R does not alter the epithelial cell invasive or seeding capacity (Fig. 4c). Furthermore, removing Wnt1 tumor epithelial cells from the primary TME allows them to establish lung metastases supporting the conclusion that intact IGF-1R signaling suppresses the ability of the Wnt1 tumor epithelial cells to leave the primary tumor. Surprisingly, macrometastases were only identified in the TVI lungs from Wnt1 and K8iKOR-Wnt1 tumor epithelial cells (Fig. 4d). H&E-stained lung sections also showed significantly smaller micrometastases in TVI lungs from DN-Wnt1 tumor epithelial cells 1 wpi (Fig. 4b).

**Figure 4.**
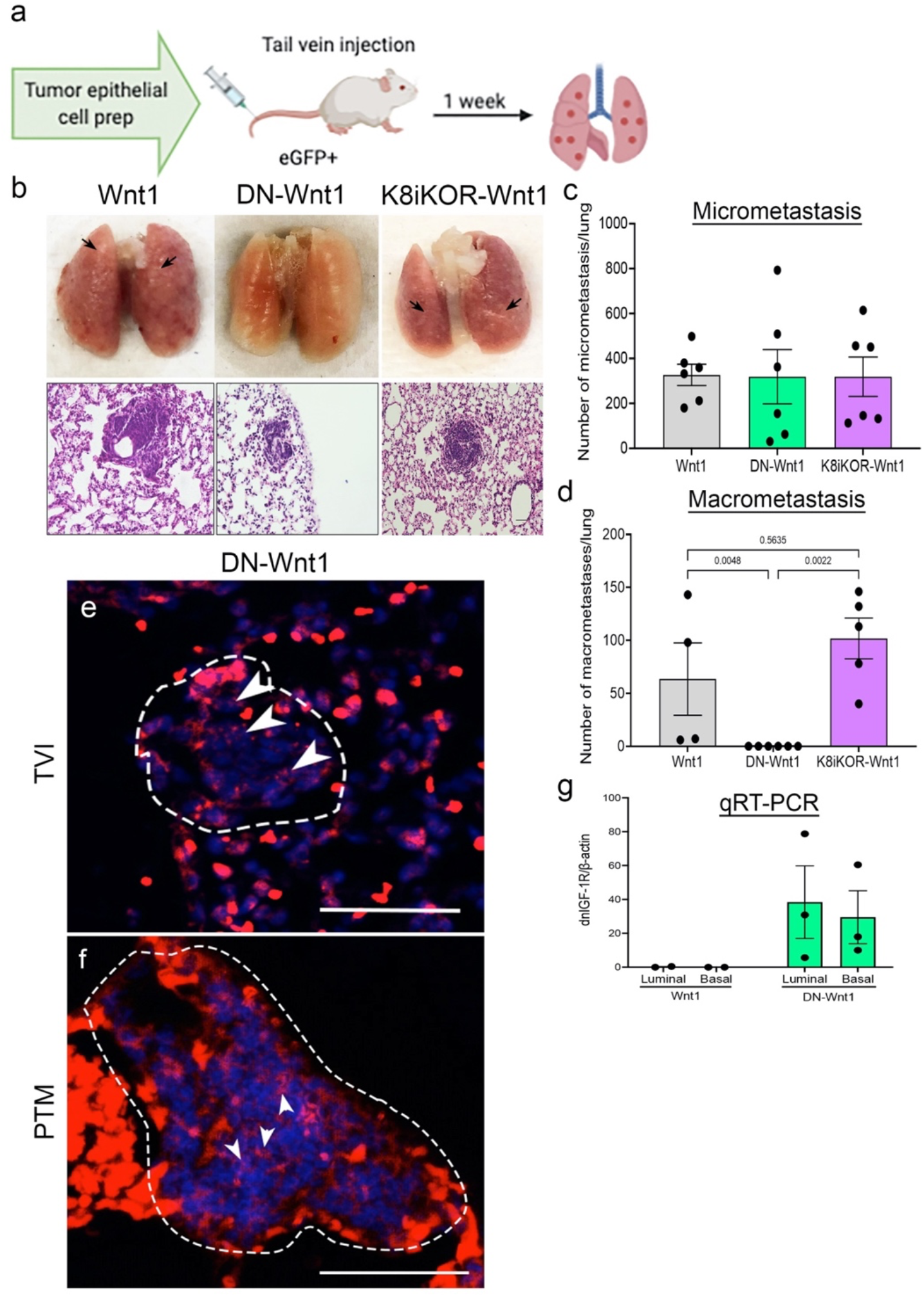
Tail vein injections of primary tumor epithelial cells with reduced IGF- 1R. **a.** Schematic of tumor epithelial cell tail vein injection. **b.** Top Row: Representative whole lung images one-week after TVI of Wnt, DN-Wnt1, or K8iKOR-Wnt1 tumor epithelial cells. Arrows denote macroscopic metastases. Bottom Row: Representative lung hematoxylin and eosin staining for micrometastases from TVI of Wnt1, DN-Wnt1, or K8iKOR-Wnt1 tumor epithelial cells. Scale bar = 50 microns. **c-d.** Micrometastases **(c)** and macrometastases **(d)** counts from TVI lungs. *Statistic:* Non-parametric Kolmogorov Smirnov test **e-f.** RNAscope immunofluorescence for the human IGF-1R transgene (*dnIGF-1R*) in DN-Wnt1 TVI micrometastases 1 wpi **(e)** or in DN-Wnt1 endogenous primary tumor micrometastases (PTM) **(f)**. Scale bar = 50 microns **g.** RT-PCR for the human IGF-1R transgene in sorted primary tumor cells from Wnt1 and DN- Wnt1 tumors.

The TVI results were surprising since the endogenous tumors that form in both the DN-Wnt1 and K8iKOR-Wnt1 models lead to a high rate of metastasis. This suggests that the process of dissociating the tumor epithelial cells alters the growth potential of the DN-Wnt1 cells once the metastases have seeded. One possible explanation for the discrepancy in lung metastatic growth after TVI between the two models is lineage specificity of the IGF-1R disruption. The MMTV promoter is active early in the mammary epithelial lineage such that both lineages express the transgene (26). RNAscope immunofluorescence analysis for the human *dnIGF-1R* transgene confirmed expression in hyperplastic mammary glands and tumors from the DN-Wnt1 mice (Supp. Fig. 5a-h) as well as in micrometastases in TVI lungs from DN-Wnt1 tumor epithelial cells 1 wpi and in endogenous primary micrometastases (Fig. 4e,f; Supp Fig. 5i-k). We then verified the expression of the *dnIGF-1R* transgene in both luminal and basal epithelial lineages by performing qRT-PCR for the human *dnIGF-1R* transgene in tumor epithelial cells following FACS (Fig. 4g). These findings support the hypothesis that the differences in the TVI metastasis phenotype from the two IGF-1R models may be due to disruption of IGF-1R in only the luminal lineage (K8iKOR-Wnt1) versus in both the luminal and basal lineages (DN-Wnt1).

### Expansion of the metastatic tumor epithelial population with reduced IGF-1R

We then asked 1) what are the cells from the DN-Wnt1 or K8iKOR-Wnt1 primary tumors that seed lung metastases, 2) what properties of the epithelial cells from the DN- Wnt1 tumor cells prevent their proliferation after TVI, and 3) why does luminal-specific deletion of *Igf1r* maintain metastatic tumor growth after TVI? To address these questions, we restricted the scRNA-Seq analysis to the tumor epithelial cell populations. Unsupervised clustering using UMAP resulted in 13 distinct epithelial populations (E0- E13) from 2,543 cells from Wnt1, DN-Wnt1, and K8iKOR-Wnt1 tumors (Fig. 5a). Using Seurat analysis for keratin expression and heat map analysis of known epithelial cell population markers (27) we identified the epithelial clusters as: alveolar (E0,E1), luminal (E3,E4,E12), differentiated luminal (E6,E10), luminal progenitor (E2) and basal (E5,E7,E8,E9,E11), one of which (E7) had high expression of the bipotential cell marker Lgr5 (Fig. 5b-c, Supp. Fig. 6; see Supp. Methods). Importantly, 3 of the 5 differentiated basal cell clusters (E5, E7, and E8) and the luminal progenitor cluster (E2) were expanded in both the K8iKOR-Wnt1 and DN-Wnt1 tumors (Fig. 5d). The expansion of the basal and luminal progenitor populations in the DN-Wnt1 and K8iKOR-Wnt1 tumors was supported by flow cytometry analyses (Supp. Fig. 7 and (23)).

**Figure 5.**
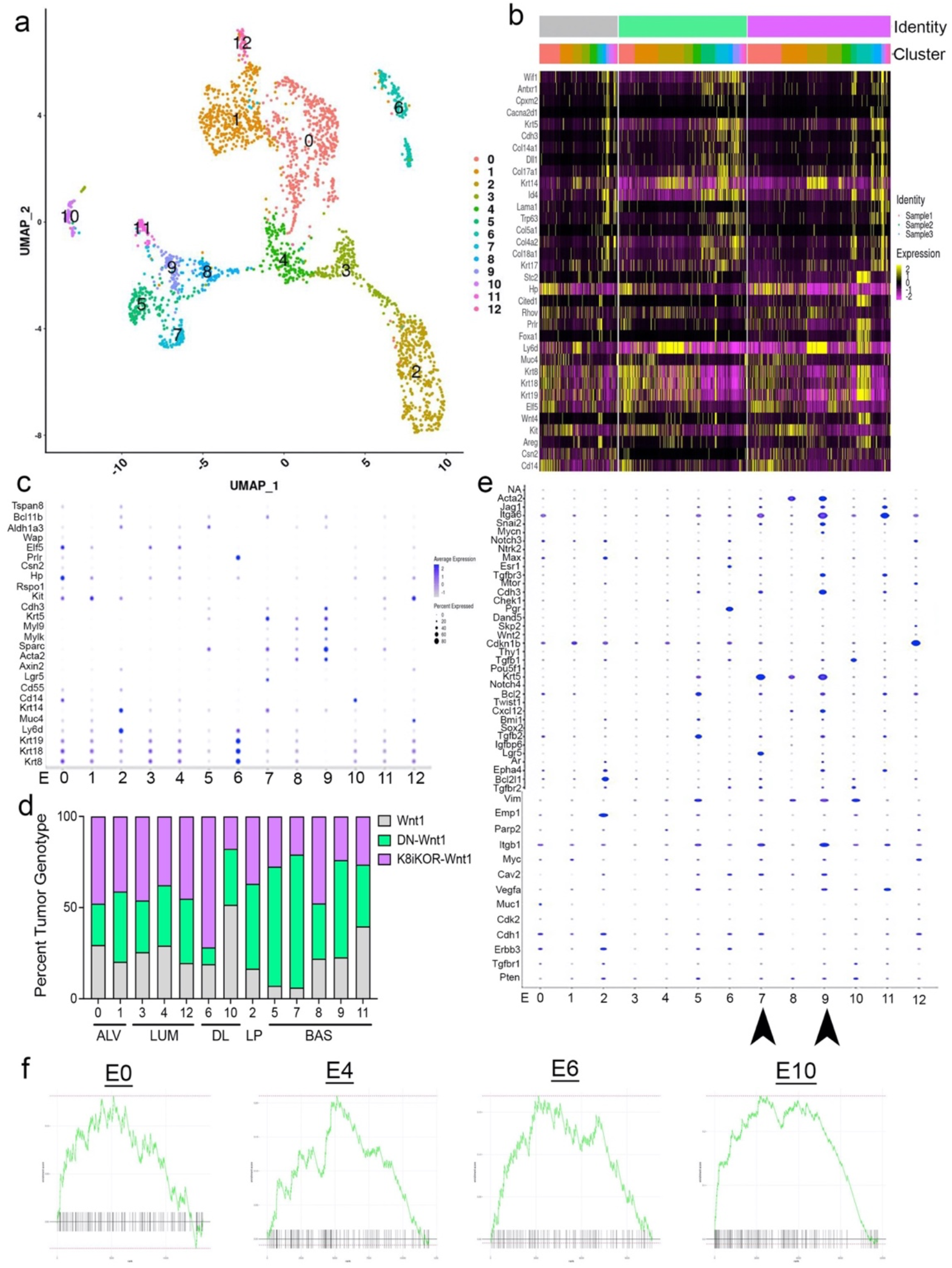
Epithelial cell populations are altered with reduced IGF-1R. **a.** UMAP plot of re-clustering of epithelial cells from Wnt1, DN-Wnt1, and K8iKOR-Wnt1 tumors resulting in 13 clusters. **b.** Heat map of top epithelial cell type markers. Top legend: top row=tumor identity: gray=Wnt1, green=DN-Wnt1, pink=K8iKOR-Wnt1; Bottom row=epithelial cell cluster. **c.** Dot plot of epithelial cell markers. **d.** Percent tumor genotype graph for each cell cluster labelled with each cell type defined by markers. (ALV=alveolar cell, LUM=luminal cell, DL=differentiated luminal cell, LP=luminal progenitor, BAS=basal cell) **e.** Dot plot of alignment with metastatic signature. Arrows depict clusters with high expression of markers indicating metastatic cell type. **f.** GSEA plots for the epithelial mesenchymal transition (EMT) hallmark signature in Clusters E0, E4, E6, and E10.

Clusters E7 and E9 are most closely linked to a previously identified metastatic signature (28) (Fig. 5e) consistent with increased metastasis in the IGF-1R deficient tumor models (Fig. 2d,e). Gene Set Enrichment Analysis (GSEA) confirmed enrichment in EMT (Fig. 5f; Supp. Fig. 8). IPA revealed key changes in cell migration and invasion pathways specific to clusters E4, E5, E8, and E11 (Supp. Fig. 9). Increased EMT and migration/invasion transcripts (Fig. 5b-e) suggests these populations are gaining mesenchymal characteristics consistent with increased metastatic potential in both basal and luminal populations in the IGF-1R deficient tumors.

### Attenuated IGF-1R decreases TVI metastatic growth by altering cell cycle

We next asked what properties of the DN-Wnt1 cells resulted in the failure of the micrometastases to proliferate and form macrometastasis in TVI mice. Analysis of lungs from TVI mice past 1 wpi revealed decreased micrometastases in DN-Wnt1 TVI lungs over time (Fig. 6b-c) suggesting maintenance of tumor epithelial cell proliferation and survival is inhibited with attenuated IGF-1R when removed from the primary tumor niche and dissociated prior to colonizing the lung. Furthermore, H&E staining revealed maintenance of micrometastases at 3 and 6 wpi but degradation at 8 and 12 wpi (Fig. 6b). In contrast, lungs from TVI mice injected with either Wnt1 or K8iKOR-Wnt1 tumor epithelial cells formed numerous macrometastases by 3 wpi (Fig. 6b).

**Figure 6.**
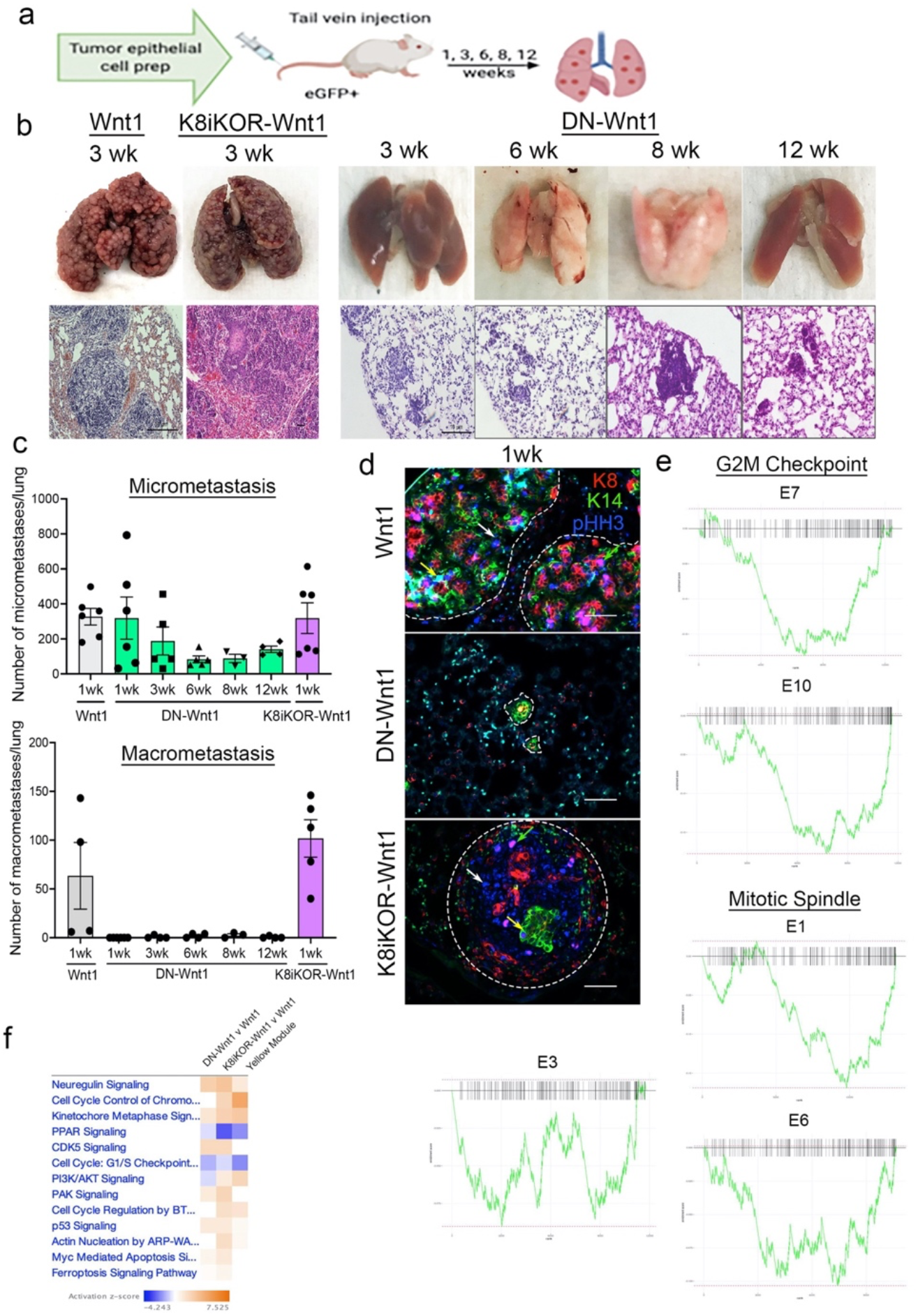
Reduced IGF-1R reduces tumor cell growth and survival in lungs after tail vein injection. **a.** Schematic of tumor epithelial cell tail vein injection. **b.** Top Row: Representative whole lung images after 3-, 6-, 8-, and 12-week TVI from Wnt1, DN- Wnt1, or K8iKOR-Wnt1 tumor epithelial cells. Bottom Row: Representative lung hematoxylin and eosin staining for micrometastases from TVI of Wnt1, DN-Wnt1, or K8iKOR-Wnt1 tumor epithelial cells over time. **c.** Micrometastases (top graph) and macrometastases (bottom graph) counts from TVI lungs over time. **d.** Immunofluorescence for K8 (red), K14 (green), and phospho-histone H3 (pHH3, blue) in metastases after 1wk TVI of Wnt1, DN-Wnt1, or K8iKOR-Wnt1 primary epithelial cells. Scale bar = 50 micron; representative of n=3. **e.** GSEA plots of G2M and Mitotic Spindle hallmark signatures in differentially expressed genes from DN-Wnt1 vs. Wnt1 epithelial cell clusters. **f.** IPA canonical pathway heat map of differentially expressed genes from DN-Wnt1 and K8iKOR-Wnt1 compared to Wnt1 tumors and the METABRIC (human) yellow (cell cycle) module.

To assess the role for IGF-1R in tumor epithelial cell proliferation we stained tissue sections for phospho-histone H3 (pHH3). Interestingly, pHH3+ expression was similar in hyperplastic glands and primary tumors from Wnt1, DN-Wnt1, and K8iKOR- Wnt1 mice (Supp. Fig. 10). In contrast, pHH3 was undetectable at 1 wpi in TVI micrometastases from DN-Wnt1 tumor epithelial cells but was detected in numerous cells in TVI micrometastases from mice injected with Wnt1 and K8iKOR-Wnt1 tumor epithelial cells (Fig. 6d). These data suggest that attenuation of IGF-1R inhibits epithelial cell proliferation when cells are dissociated after removal from the primary tumor niche. However, signature analysis by GSEA showed reduced enrichment for genes associated with G2M checkpoint in clusters E7 and E10 and for genes associated with mitotic spindle in clusters E1, E3 and E6 of the DN-Wnt1 vs Wnt1 tumor cells (Fig. 6e) supporting alterations in proliferation in specific populations in the primary tumor.

Comparison of the METABRIC cell cycle signature (Fig. 1b) with the whole mouse tumor scRNA-seq transcripts revealed correlation of several cell cycle pathways including cell cycle checkpoint, chromosome regulation, and apoptosis signaling further supporting that loss of IGF-1R alters the cell cycle through changes in gene expression (Fig. 6f).

### Collective metastatic seeding is diminished in tumor epithelial cells in DN-Wnt1 model

The lack of proliferation in the DN-Wnt1 TVI metastases explains the observed phenotype but does not explain the underlying reason for TVI metastases maintenance and growth formed from K8iKOR-Wnt1 tumor cells. In addition to cell cycle changes in the tumor epithelial cells with reduced IGF-1R, we also observe changes in pathways involved in cell adhesion (Fig. 7). Several studies have reported collective cell invasion is necessary to seed metastatic lesions and promote metastatic growth (29–31). This invasion is dependent on K14^+^ leader cells adhered to clusters of epithelial cells to initiate collective invasion (29, 32). Micrometastases from TVI mice with either Wnt1 or K8iKOR-Wnt1 tumor epithelial cells at 1 wpi were composed of K14^+^ basal cells, K8^+^ luminal cells, and a few K8^+^/K14^+^ double-positive cells (Fig. 7a-c). In contrast, lungs from DN-Wnt1 TVI mice at 1 wpi were composed mostly of K14^+^ leader cells (Fig. 7a-c). These results suggest loss of IGF-1R expands the K14^+^ leader cell population, but phenotypic alterations within these cells decrease collective invasion to reduce metastatic proliferation. One interesting question is which cell population(s) contribute to the K14^+^ leader cells. Previous reports have shown K8^+^ luminal cells gain K14 expression and can also participate as metastatic, leader cells (29, 32). Both the DN- Wnt1 and K8iKOR-Wnt1 tumors have increased basal cell populations (Fig. 5d); however, we also found that sorted luminal cell populations from both tumors showed increased expression of K14 compared to Wnt1 tumors (Fig. 7d). Thus, both basal and luminal populations in tumors with reduced IGF-1R could be contributing K14^+^ leader cells to seed metastases.

**Figure 7.**
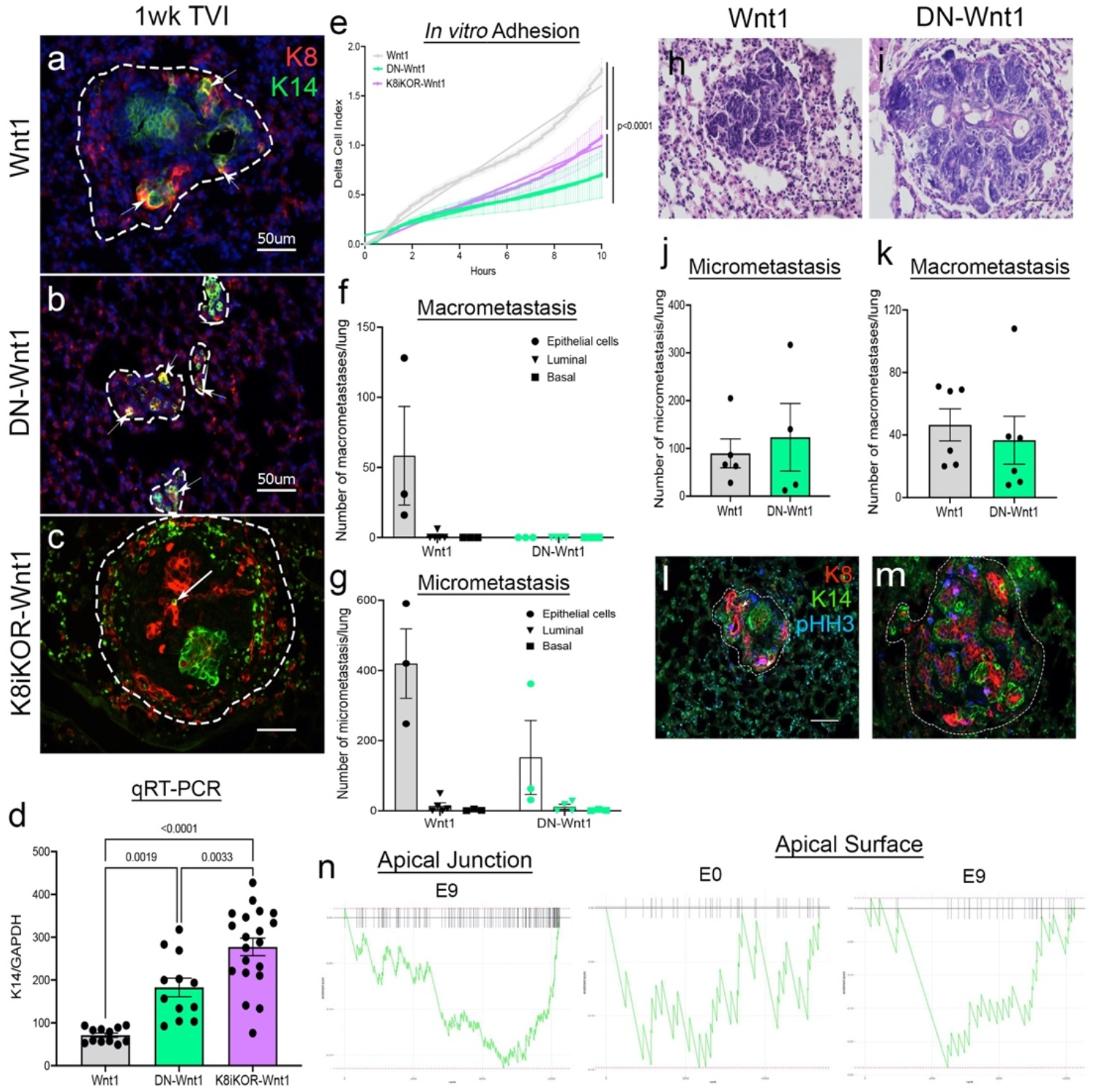
Reduced IGF-1R function decreases tumor cell adhesion. **a-c.** Representative TVI lung staining for K8 (red) and K14 (green) from Wnt1 **(a)**, DN-Wnt1 **(b)** and K8iKOR-Wnt1 **(c)** 1 wpi mice. Dotted lines demarcate the metastatic lesions. Arrows indicate double positive K8 and K14 cells. scale bar=50 micron; n=3. **d.** RT-PCR for K14 in sorted luminal (CD24^+^/CD29^lo^) primary tumor cells from Wnt1, DN-Wnt1, and K8iKOR-Wnt1 tumors showing increased expression in both IGF-1R models. *Statistic:* Mann-Whitney U non-parametric t test. **e.** Measurement of adhesion from Wnt1 (grey), DN-Wnt1 (green), or K8iKOR-Wnt1 (purple) by delta cell index over time for 6 hours using the real-time xCELLigence assay. n=3; *Statistic:* Non-linear regression least squares regression for slope best fit p<0.0001 for Wnt1 control compared to DN-Wnt1 or K8iKOR-Wnt1 **f-g.** Macrometastases **(f)** or micrometastases **(g)** counts in lungs from TVI mice injected with Wnt1 or DN-Wnt1 tumor epithelial cells (250K), sorted luminal epithelial cells, or sorted basal epithelial cells after 1 wpi. **h-i.** Representative H&E images from Wnt1 **(h)** and DN-Wnt1 **(i)** lung micrometastases from TVI mice injected with Wnt1 or DN-Wnt1 tumor epithelial cells cultured overnight at low adherence. **j-k.** Micrometastases **(j)** and macrometastases **(k)** counts in lungs from TVI mice injected with Wnt1 or DN-Wnt1 tumor epithelial cells cultured overnight at low adherence. **l-m.** Immunofluorescence of K8 (red), K14 (green), and pHH3 (blue) in Wnt1 **(l)** and DN- Wnt1 **(m)** TVI micrometastases from tumor epithelial cells cultured overnight at low adherence. **n.** GSEA plots for the apical junction and apical surface hallmark signatures from differentially expressed genes in DN-Wnt1 vs. Wnt1 epithelial tumor cell clusters.

### Cell adherence is altered in tumor epithelial cells with decreased IGF-1R function

Smaller micrometastases composed of mostly K14^+^ cells observed in the DN- Wnt1 TVI lungs suggest a defect in adherence between the K14^+^ leader cells and the metastatic proliferating cells. Adherence was decreased in DN-Wnt1 and K8iKOR-Wnt1 compared to Wnt1 primary tumor epithelial cells in vitro (Fig. 7e). Consistent with these findings, the number of DN-Wnt1 primary tumor epithelial cell clusters and single tumor epithelial cells had decreased adherence to collagen matrix compared to Wnt1 primary tumor cells (Supp. Fig. 11a-e). In contrast, there was no significant difference between the K8iKOR-Wnt1 and Wnt1 primary tumor epithelial cells in their ability to adhere to collagen (Supp. Fig. 11a-e). Immunofluorescence revealed increased K14^+^ and decreased K8+ cell adherence from DN-Wnt1 compared to Wnt1 primary tumors both in clusters and individual cells (Supp. Fig.11f,g) consistent with *in vivo* TVI analysis (Fig. 7a-c). Moreover, the non-adherent cells from the DN-Wnt1 tumors had increased E-cadherin and cyclin D1 expression indicating these are the proliferating luminal epithelial cells (Supp. Fig. 11h,i). Taken together, these data support the conclusion that attenuated IGF-1R with the *dnIGF-1R* transgene alters adherence between the K14^+^ leader cell and other epithelial cells, particularly those that are necessary to proliferate in the metastatic lesion, while deletion of luminal *Igf1r* does not alter adherence to the same extent. These findings support the hypothesis that disruption of IGF-1R in both the luminal and basal lineages in the DN-Wnt1 tumors (Fig. 4) is necessary to disrupt adhesion between epithelial cells.

Our data and other recent studies (33–36) indicate that cell adherence is necessary to promote metastatic seeding and growth. We next asked whether paracrine signaling between the epithelial cell populations is necessary to maintain metastatic growth. Primary tumor epithelial cells from Wnt1 or DN-Wnt1 mice were sorted for luminal (CD24^+^/CD29^lo^) and basal (CD24^+^/CD29^hi^) epithelial populations and injected into the tail vein of eGFP mice. Importantly, lung macrometastases were significantly decreased in the sorted populations from Wnt1 tumor compared to combined epithelial cell TVIs (Fig. 7f). Similarly, we observed decreased lung micrometastases from both Wnt1 and DN-Wnt1 sorted population TVIs compared to combined epithelial cells (Fig. 7g). These data support the conclusion that luminal and basal epithelial cell adherence is necessary for metastatic seeding and growth.

To test whether restoration of lineage associations would restore metastatic growth in the DN-Wnt1 tumor cells, we performed TVIs with primary epithelial cells from Wnt1 or DN-Wnt1 tumors cultured overnight in low adherent plates to allow for cell re-adherence after dissociation. Enhancing cell adhesion by overnight incubation resulted in similar numbers of macrometastases and micrometastases in DN-Wnt1 and Wnt1 TVI primary tumor epithelial cells (Fig. 7h-k). Further analysis by immunofluorescence revealed pHH3^+^ and K8^+^ micrometastases from DN-Wnt1 epithelial cells 1 wpi similar to Wnt1 micrometastases (Fig. 5, Fig. 7l,m) suggesting decreased cell adherence in DN- Wnt1 tumor epithelial cells is amplified by tumor dissociation. Analysis of adhesion target transcripts identified upregulation of adhesion in Cluster E2 (luminal progenitor) across all tumor types and downregulation of adhesion in the basal cell clusters (Clusters E5, E7, E8, and E9) in DN-Wnt1 tumors (Fig. 7n, Supp. Fig 12a).

### Cell adherence is dysregulated by enhanced P-cadherin expression in epithelial cells with reduced IGF-1R function

Recently, the Ewald lab reported E-cadherin loss is required for metastatic invasion, and its re-expression is necessary to promote metastatic growth (37). Furthermore, upregulation of P-cadherin and its co-expression with E-cadherin in the primary tumor is a marker of more aggressive, metastatic breast tumors (38–41). To determine if cadherin expression changes with IGF-1R expression in patient tumors, we analyzed the METABRIC dataset and identified an inverse correlation of P-cadherin and IGF-1R expression. Conversely, E-cadherin is positively correlated with IGF-1R expression across all breast tumors (Fig. 8a).

To determine whether cadherin expression is similarly altered in the DN-Wnt1 tumors, we screened for cadherin expression in each epithelial cluster from the scRNA-Seq data. As expected, luminal cell types had higher E-cadherin (Cdh1) expression whereas basal cell types had higher P-cadherin (Cdh3) and T-cadherin (Cdh13) expression (Supp. Fig. 12b). E-cadherin expression in DN-Wnt1 tumor epithelial cells by qRT-PCR was decreased compared to Wnt1 cells (Supp. Fig. 12c). Moreover, E-cadherin levels were decreased in clusters E5 and E7 (Fig. 8b), which also expressed P-cadherin (Fig. 8c). Immunostaining similarly showed decreased E-cadherin and increased P-cadherin protein expression in DN-Wnt1 and K8iKOR-Wnt1 primary tumors compared to Wnt1 tumors (Fig. 8d-m). Interestingly, total E-cadherin expression was altered primarily at the protein level, while P-cadherin was changed at both the RNA and protein levels in tumors with reduced IGF-1R. Importantly, co-expression of E-cadherin and P-cadherin was increased in DN-Wnt1 and K8iKOR-Wnt1 tumors (Fig. 8d- m). Thus, reduced IGF-1R was associated with altered E-cadherin and P-cadherin in tumor epithelial cells.

**Figure 8.**
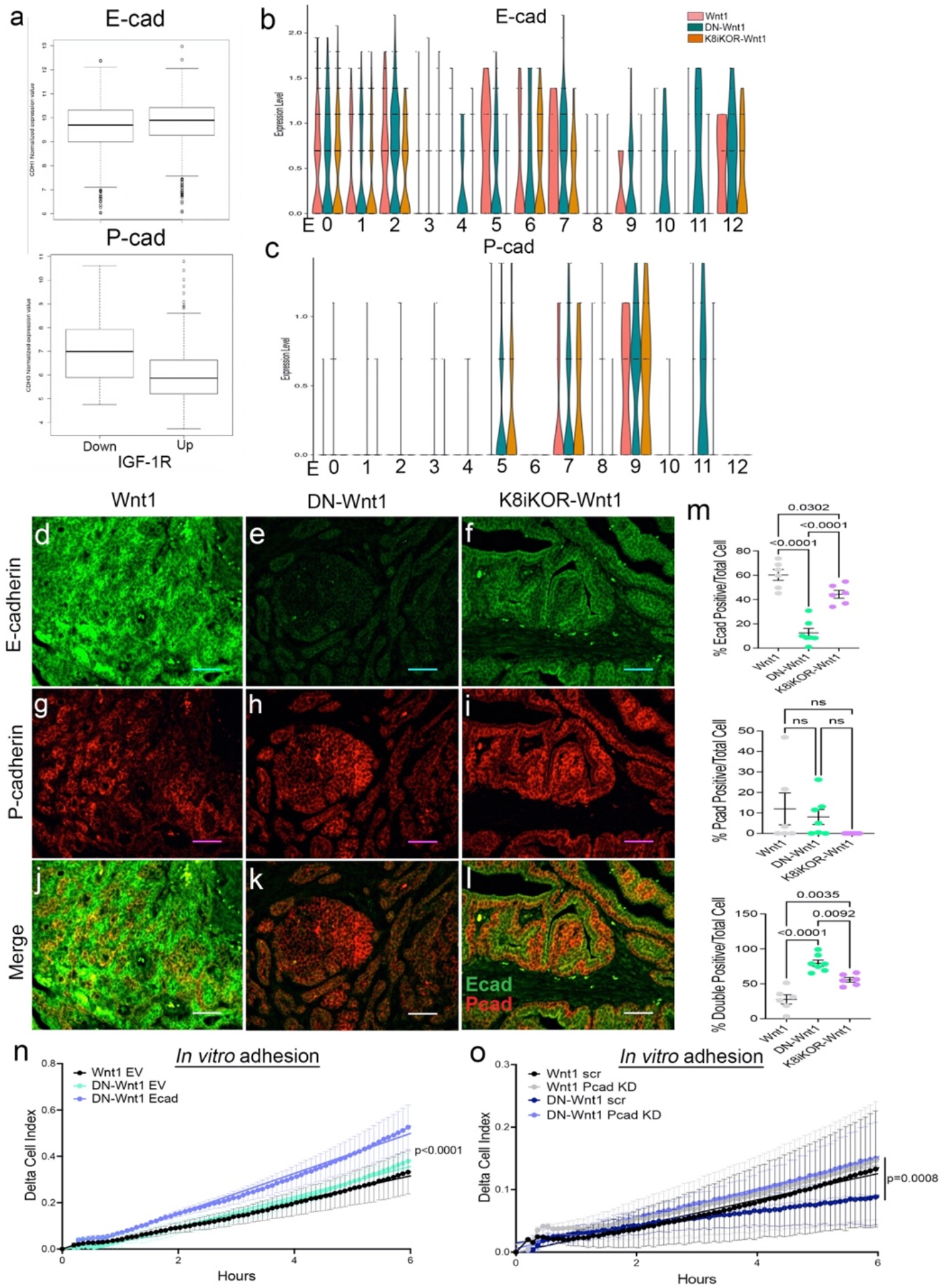
Altered cadherin expression in tumors with reduced IGF-1R. **a.** METABRIC data analysis for E-cadherin or P-cadherin in patient tumors with low IGF- 1R (IGF-1R z-score < -1) or high IGF-1R (IGF-1R z-score > 1) (p < 2.0×10^-16^) *Statistic:* Student’s *t*-test. **b-c.** E-cadherin **(b)** and P-cadherin **(c)** expression in each epithelial cell cluster identified with single-cell sequencing in Wnt1, DN-Wnt1, or K8iKOR-Wnt1 primary tumors. **d-m.** Representative images of E-cadherin (red) or P-cadherin (green) immunostaining in Wnt1 **(d, g, j)**, DN-Wnt1 **(e, h, k)**, and K8iKOR-Wnt1 **(f, i, l)** primary tumors. **m.** E-cadherin, P-cadherin, and double positive cell count graphs of primary tumors. *Statistic:* One-Way ANOVA with Tukey’s Multiple Comparison post-hoc test **n.** Adhesion (delta cell index) over time in Wnt1 or DN-Wnt1 primary tumors with empty vector (EV) or E-cadherin overexpression (Ecad). n=3; *Statistic:* Non-linear regression. **o.** Adhesion (delta cell index) over time in Wnt1 or DN-Wnt1 with P-cadherin knockdown (Pcad KD). n=3; *Statistic:* Non-linear regression.

To test the functional role of altered E-cadherin and P-cadherin in cells with attenuated IGF-1R, we first transiently re-expressed E-cadherin in DN-Wnt1 primary tumor epithelial cells and measured cell adhesion *in vitro*. Overexpression of E-cadherin increased epithelial cell adhesion compared to empty vector control (Fig. 8n). Furthermore, reducing P-cadherin in DNA-Wnt1 primary tumor epithelial cells significantly increased tumor adhesion restoring adhesion back to the level of the Wnt1 tumor cells (Fig. 8o). Thus, altering cadherins in DN-Wnt1 primary tumor epithelial cells rescues the compromised adherence suggesting these changes in E- and P-cadherins due to reduced IGF-1R are necessary for metastasis.

## Discussion

A major question in cancer biology is how do primary tumor cells metastasize to another site? Here we show loss of IGF-1R in the primary tumor promotes metastasis by modulating cadherin expression, altering epithelial cell properties and increasing basal leader cells for collective invasion. Furthermore, reduced IGF-1R function or expression increases metastatic extravasation. Surprisingly, once the primary tumor cells are removed from the tumor niche and dissociated we discovered loss of IGF-1R function promotes tumor cell quiescence.

While it is well established that epithelial cells gain mesenchymal cell properties to migrate out of the primary tumor (29–32, 36), several recent studies have shown only a subset of mesenchymal properties are necessary for migration and invasion referred to as partial EMT (42–45). While original dogma was that the metastatic process occurs by single tumor epithelial cell migration and invasion, recent observations of collective epithelial cell migration have presented a new mechanism for metastasis that relies on interactions between a mesenchymal-like leader cell with other epithelial cells in the primary tumor (36). Thus, understanding how cell-cell interactions are regulated both in the primary tumor and at distant sites of colonization is critical to determining metastatic potential of tumor cells.

Loss of E-cadherin is a hallmark of EMT and necessary for basal cells to adapt to becoming leader metastatic cells (29). The Ewald lab previously described a process by which the transition of E-cadherin expression is critical for collective invasion (37). Here, we have shown E-cadherin expression is decreased in mouse models with reduced function or expression of IGF-1R to drive collective invasion. In the TVI models, we show that loss of adhesion between epithelial cells compromises collective invasion to promote growth of the metastatic lesions. Interestingly, recent reports have shown acquisition of P-cadherin is necessary for tumor cells to become metastatic. More importantly, the co-expression of P-cadherin and E-cadherin is critical for enhanced metastasis and suggests these cells are exhibiting a partial EMT phenotype. Attenuation or reduced IGF-1R levels in the Wnt1 mouse tumor model results in co-expression of P-cadherin and E-cadherin and a partial EMT phenotype suggesting increased metastatic properties of these tumor cells.

While loss of IGF-1R is sufficient to drive a partial EMT phenotype and collective invasion to promote metastasis, alterations in the tumor microenvironment may also be required for increased tumor extravasation. Our previous studies showed heightened cell stress driven by attenuated IGF-1R resulted in immune cell evasion and a pro-metastatic tumor microenvironment (14). Furthermore, the TVI model demonstrated that removing the Wnt1 tumor cells from their primary microenvironment was sufficient to promote their metastasis after TVI suggesting alterations in the TME driven by attenuating IGF-1R promote metastasis.

While a similar metastatic process is observed in the DN-Wnt1 and K8iKOR- Wnt1 primary tumor models, the TVI experiments revealed clear differences in the phenotype of the primary tumor cells in these models. There are two key differences in these models that likely contribute to these findings: 1) the DN-Wnt1 model attenuates the receptor activity whereas the K8iKOR-Wnt1 model is a gene knockout in the luminal epithelium, and 2) the *dnIGF-1R* transgene is expressed in luminal and basal epithelial cells blocking the receptor function in all mammary epithelium, whereas receptor expression is decreased only in the luminal epithelial cells in the K8iKOR-Wnt1 model leaving the basal cell IGF-1R intact. Potentially, the loss of IGF-1R function in both luminal and basal epithelial cells may lead to the observed TVI model phenotype because of reduced adherence. These findings emphasize modeling importance.

It is clear from the spontaneous tumor models attenuated or loss of IGF-1R decreases tumor latency and increases metastasis. These results are consistent with the clinical data where trials inhibiting IGF-1R have been unsuccessful. The interconnectedness of the tumor epithelium and microenvironment is highly complex. The advantage of our models is the ability to study stochastic tumor progression in the context of the microenvironment which reveals this complex tumor biology. Importantly, the mouse modeling data aligns with the human gene expression and pathway analyses and provides a basis for understanding why loss of IGF-1R in human breast cancers is associated with a worse outcome.

## Data Availability Statement

The data generated in this study will be made publicly available upon publication.

## Supporting information

Obr Supp Figures revised

Obr Supp Methods

## Acknowledgements

This work was supported by Public Health Service National Institutes of Health grants NCI R01CA204312 (T.L.W) and NCI R01CA128799 (D.L.), New Jersey Commission on Cancer Research Postdoctoral Fellowship DFHS15PPC039 and American Cancer Society-Fairfield County Roast Postdoctoral Fellowship 130455-PF-17-244-01-CSM (A.E.O.). We thank Dr. Sukhwinder Singh of the NJMS Flow Cytometry and Immunology Core Laboratory for the assistance with flow cytometry analysis and sorting, the Office of Advanced Research Computing (OARC) at Rutgers University under NIH 1S10OD012346-01A1 for the critical work made possible through access to the Perceval Linux cluster, BioRender.com for access to create the graphical abstract, and Dr. Yi Li for providing the *MMTV-Wnt1* mice.

## Author contributions

AEO performed the majority of the experiments and statistical analyses, participated in the study design and wrote the manuscript. Y-JC performed the WGCNA METABRIC analysis. VC performed the initial analyses on the K8iKOR-Wnt1 mouse tumor line. AL performed the scRNA-Seq analyses. KM performed metastases quantification, RNAScope and participated in the *in vitro* adhesion assays and study design. JJB performed metastases quantification, qRT-PCR for *Igf1r* deletion in sorted cell populations and participated in the *in vitro* adhesion assays and study design. QS performed mouse genotyping, tamoxifen tests, gland analyses and tumor harvesting. EG and DL contributed to results interpretation and manuscript editing. TLW is the principal investigator for this project and was involved in study design, data analysis, manuscript editing and submission. All authors read and approved the final manuscript.

