## Supplementary material for "Breast tumor Insulin-like growth factor receptor regulates cell adhesion and metastasis: Alignment of mouse single cell and human breast cancer transcriptomics": Obr Supp Figures revised

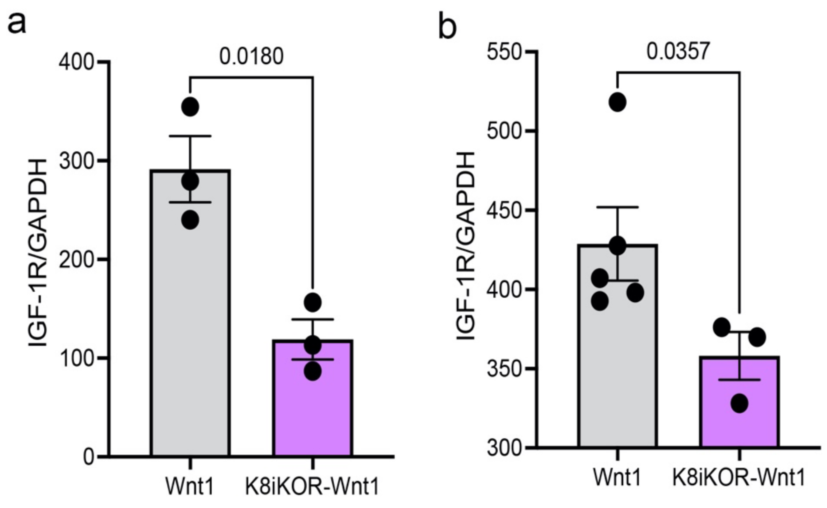


**Supplemental Figure 1: K8iKOR model validation: *Igf1r* expression in mammary tissue and tumors of K8iKOR-Wnt1 mice. a-b.** IGF-1R expression in 16-week mammary gland hyperplasias **(a)** or end stage tumors **(b)** from Wnt1 or K8iKOR-Wnt1 mice by qRT-PCR**.** *Statistic:* Unpaired parametric Welch's *t* test (hyperplasia), Unpaired non-parametric Mann-Whitney U test.

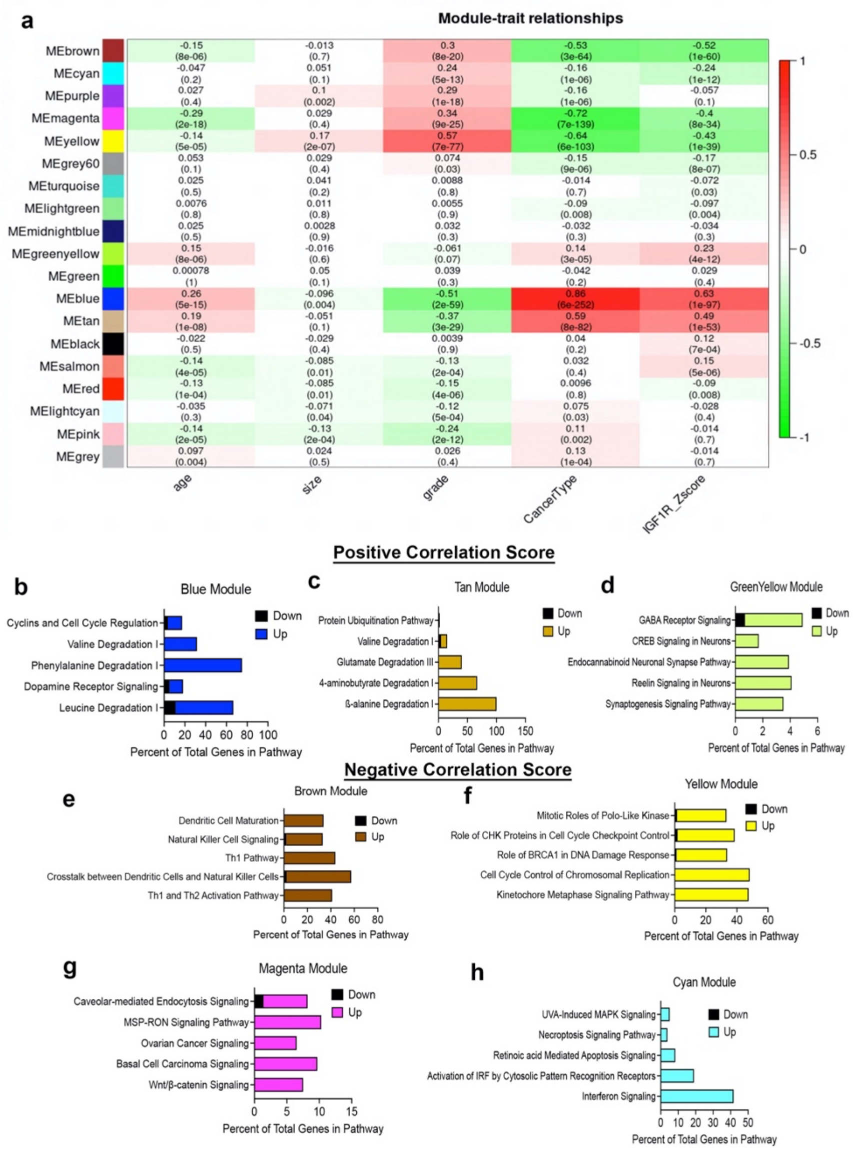


**Supplemental Figure 2: METABRIC weighted gene co-expression network analysis (WGCNA) for all gene probes associated with low or high IGF-1R expression. a.** Table of integrated WGCNA (IGF1R-GS1) showing module and clinical trait association. Each row corresponds to a module by its eigengene (ME), each column to a clinical measurement. Each cell contains the corresponding correlation and p-value (in parentheses). The table is color-coded by correlation according to the color legend. Green < 0 for negative correlation; Red > 0, for positive correlation. Initial analyses revealed three gene co-expression modules significantly correlated with high IGF-1R Z-scores (correlation score > 0.2; Supp. Fig. 2a-d) and four gene modules significantly correlated with low IGF-1R Z-scores (correlation score < -0.25; Supp. Fig. 2a, e-h). **b-d.** Top 5 pathways identified by ingenuity pathway analysis (IPA) revealing functional signatures in 3 modules positively correlated with IGF-1R expression. (blue and tan=metabolic signatures, greenyellow=neuron signature) **e-h.** Top 5 pathways identified by IPA revealing key signatures in 4 modules inversely (negative) correlated with IGF-1R expression. (brown module=immune signature, yellow module=cell cycle signature, magenta=Wnt signaling signature, cyan=apoptosis signature).

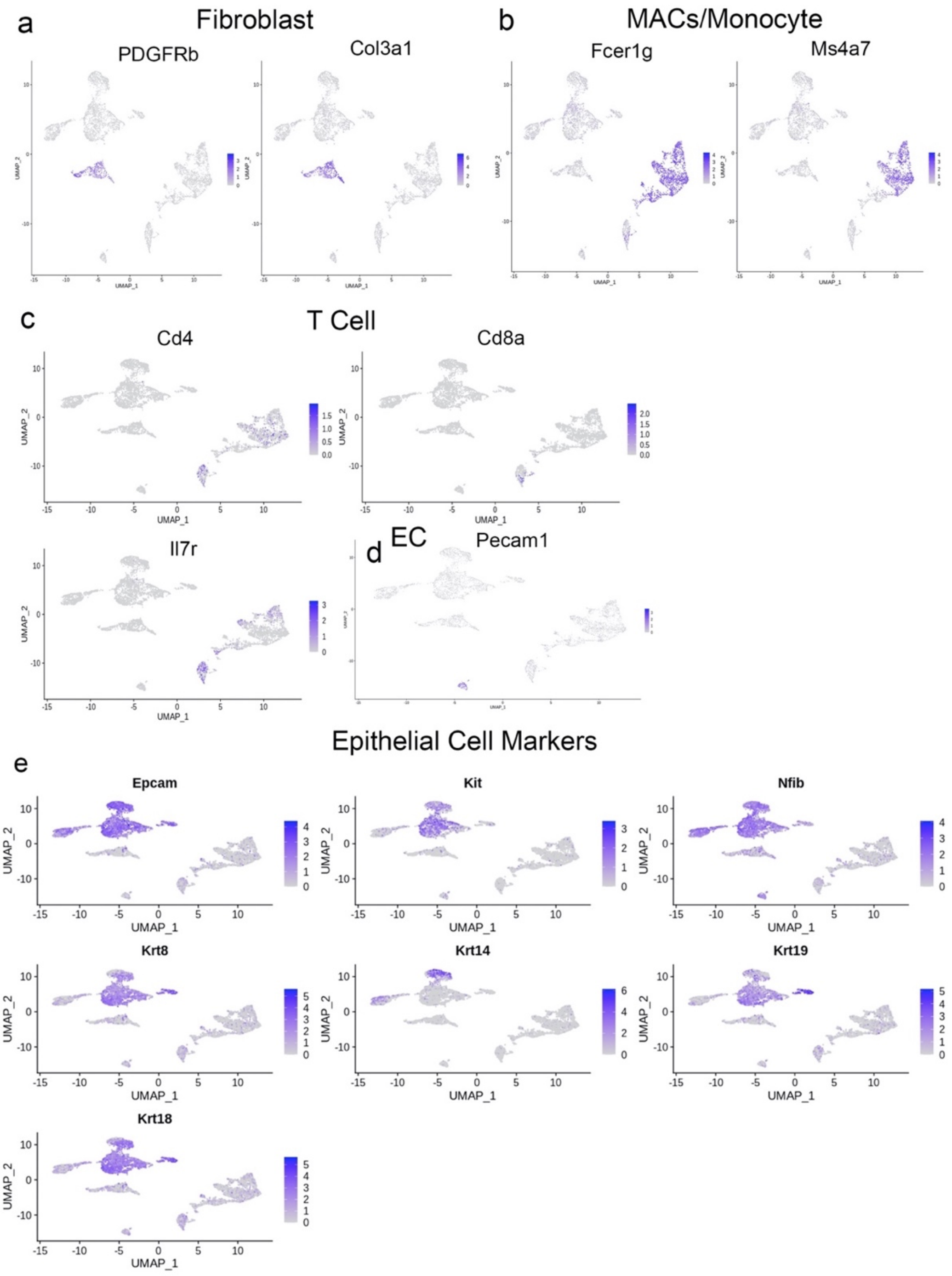


**Supplemental Figure 3: Identification of tumor cell populations.**
**a-e.** Seurat analysis plots with cell identification markers for fibroblasts (PDGFRb, Col3a1) **(a)**, MACs/monocytes (Fcer1g, Ms4a7) **(b)**, T cells (Cd4, Cd8, Il17r) **(c)**, endothelial cells (EC; Pecam1) **(d)**, and epithelial cells (Epcam, Kit, Nfib, Krt8, Krt14, Krt19, Krt18) **(e)**.

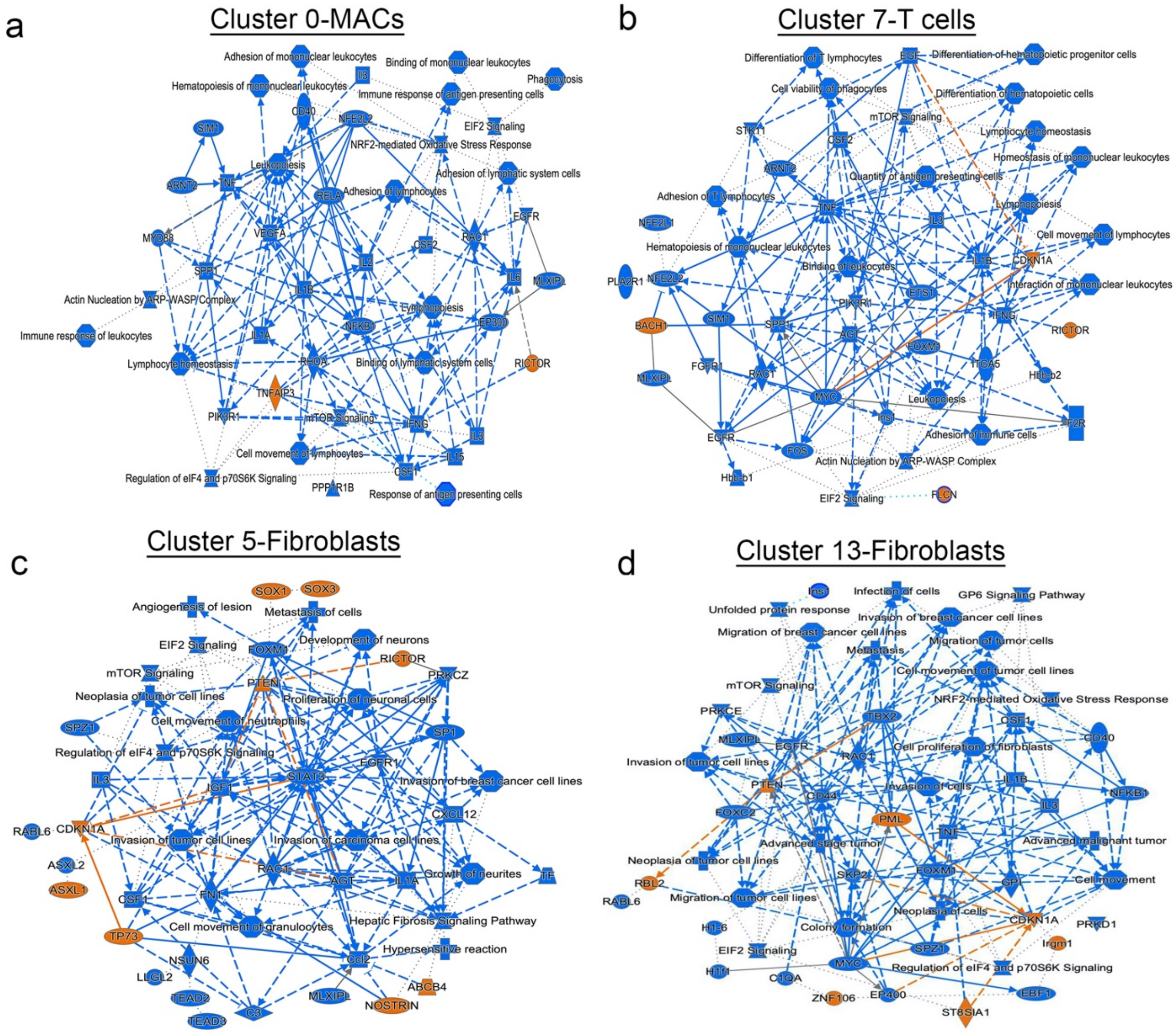


**Supplemental Figure 4: Pathway and cell function changes in tumor cells with reduced IGF-1R. a-d.** Graphical summary plots for K8iKOR-Wnt1 vs. Wnt1 tumors for Cluster 0 (MACs) **(a)**, Cluster 7 (T-cells) **(b)**, and Clusters 5 and 13 (fibroblasts) **(c,d)**.

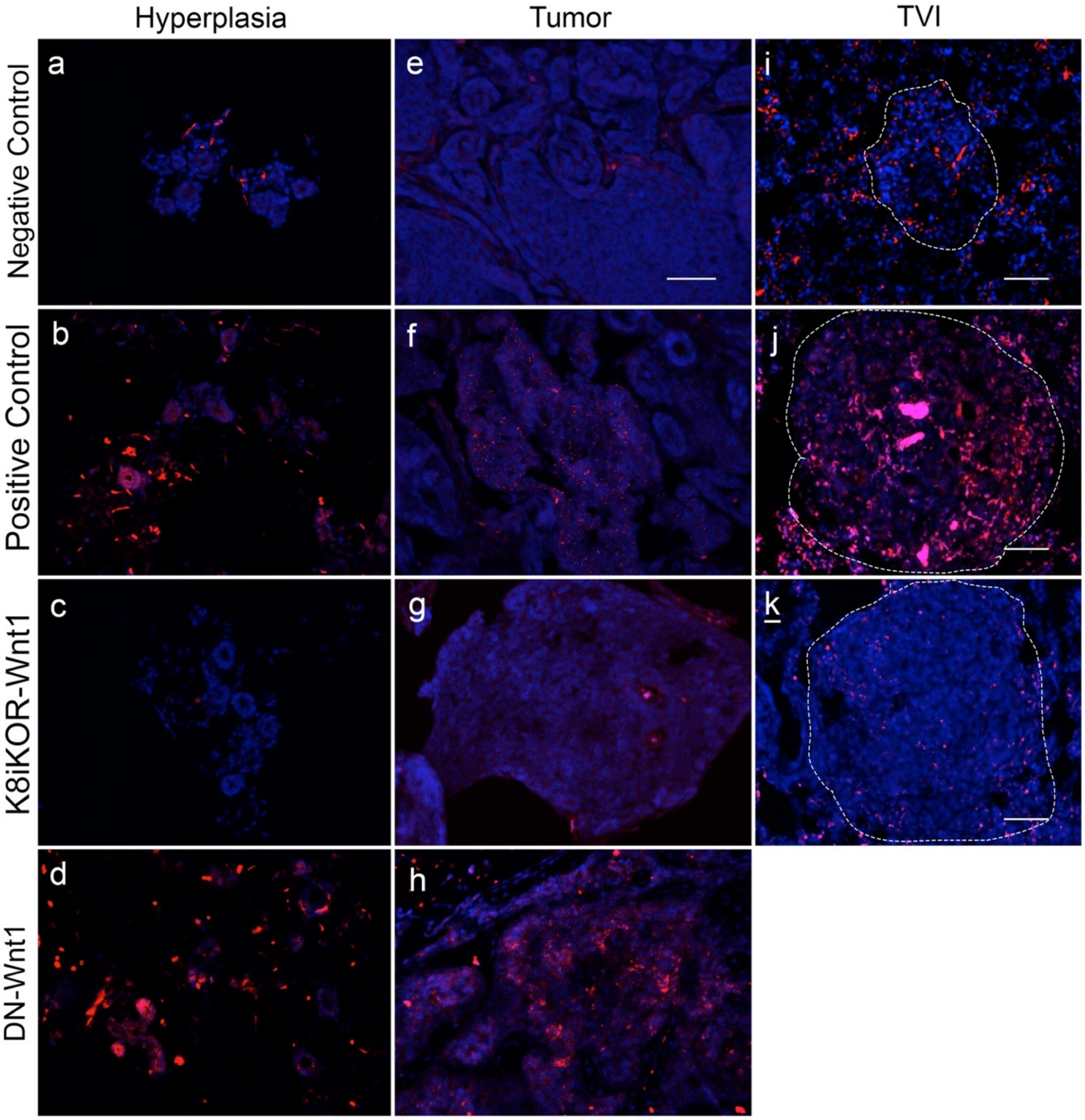


**Supplemental Figure 5: The *dnIGF-1R* transgene is expressed in luminal and basal epithelial cells in hyperplastic mammary gland and tumor tissue. a-d.** RNAscope representative images in hyperplastic mammary glands for a negative probe control **(a)**, positive housekeeping gene probe control **(b)**, and human *Igf1r* probe (dominant-negative *Igf1r* transgene) in negative K8iKOR-Wnt1 **(c)** and positive DN-Wnt1 **(d)** tissue. **e-h.** RNAscope representative images in mammary tumors as for negative probe control **(e)**, positive housekeeping gene probe control **(f)**, and human *Igf1r* probe (dominant-negative *Igf1r* transgene) in negative K8iKOR-Wnt1 **(g)** and positive DN-Wnt1 **(h)** tissue. **i-k.** RNAscope representative images in micrometastases from tail vein injection (TVI) lungs for a negative probe **(i)**, positive probe **(j)**, and human *Igf1r* probe (dominant-negative *Igf1r* transgene) in negative K8iKOR-Wnt1 tissue **(k)**. Sections are representative of 4 different hyperplasias, tumors, or TVI mets per group, scale bar=50 μm.

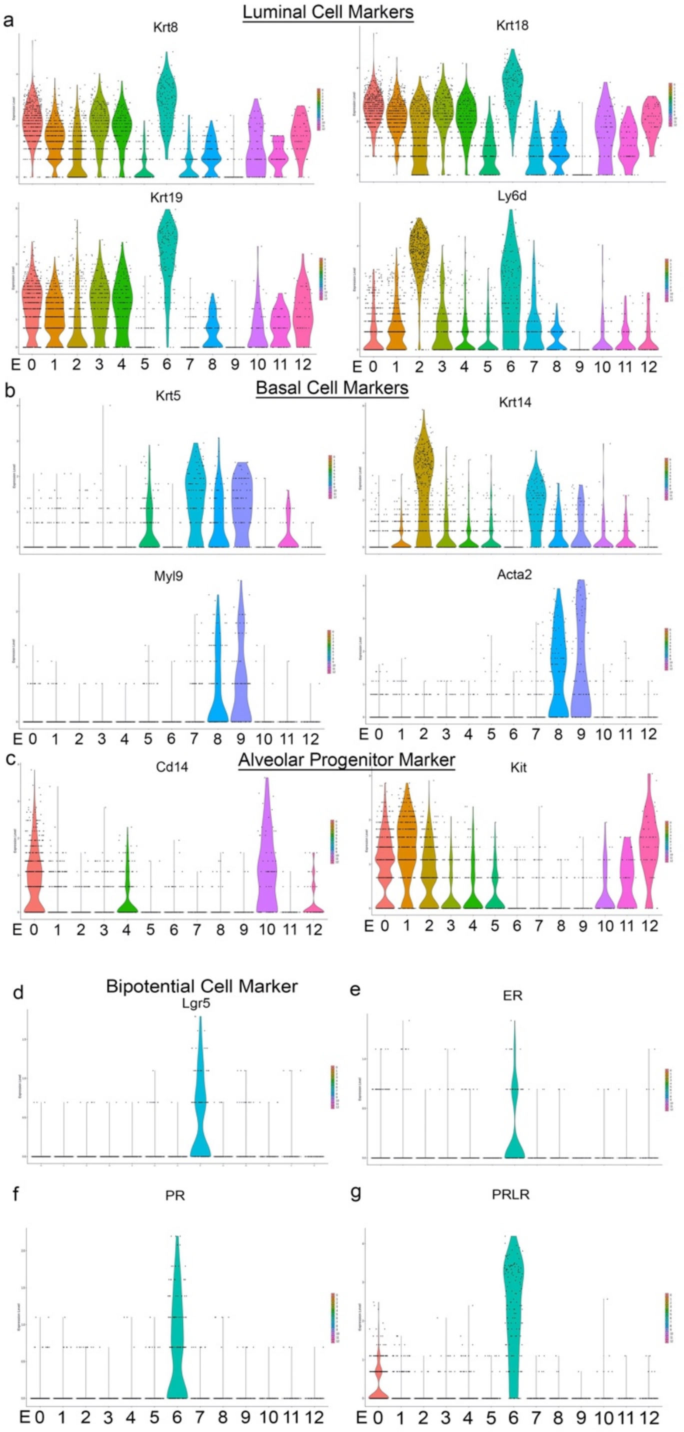


**Supplemental Figure 6: Epithelial cell subtypes in mammary tumors. a-d.** Violin plot for markers for luminal **(a)**, basal **(b)**, and alveolar cells **(c)** and bipotential progenitors **(d). e-g.** Cluster 6 has increased levels of estrogen receptor (ER) **(e)**, progesterone receptor (PR) **(f)**, prolactin receptor (PRLR) **(g)** identifying this cluster as a hormone-sensing cell.

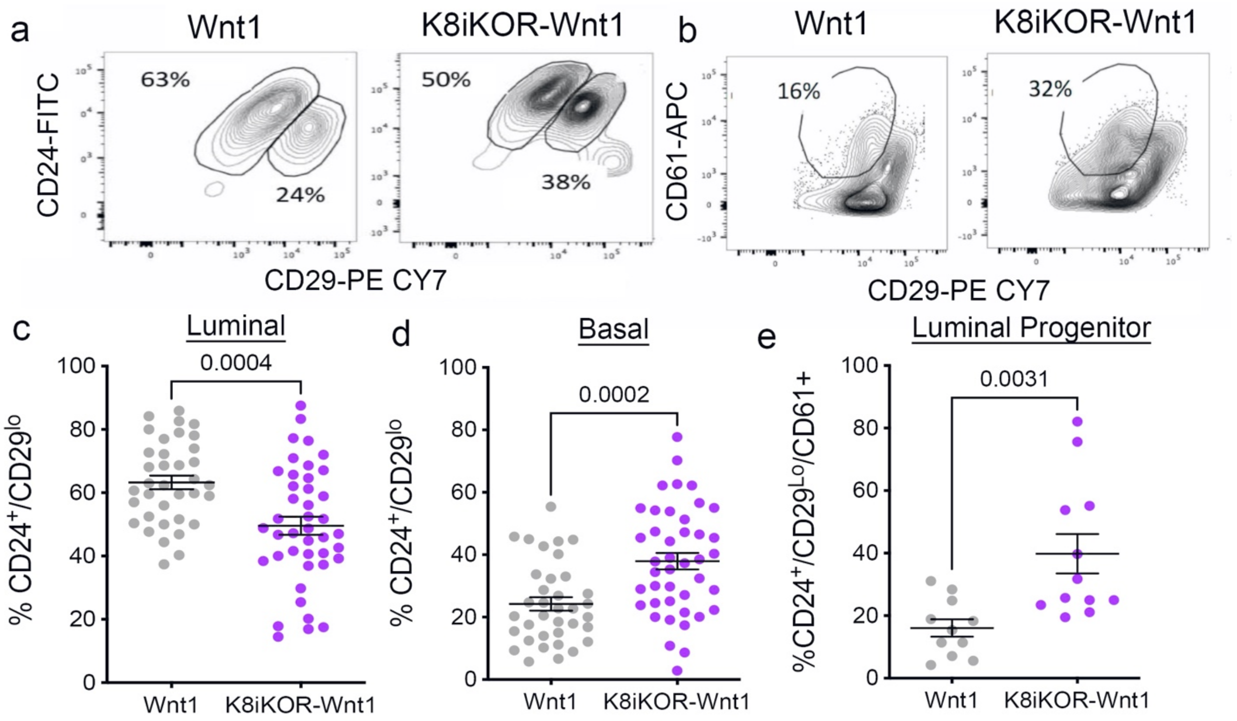


**Supplemental Figure 7: Validation of epithelial lineage changes in K8iKOR-Wnt1 tumors by flow cytometry. a-b.** Representative contour plots of flow cytometry of the $CD24^{+}/CD29^{lo}$ (luminal) and $CD24^{+}/CD29^{hi}$ (basal) cell populations **(a)** and $CD24^{+}/CD29^{lo}/CD61^{-}$ (luminal progenitor) cell population **(b)** in Wnt1 and K8iKOR-Wnt1 tumors. **c-e.** Quantification of luminal **(c)**, basal **(d)**, and luminal progenitor **(e)** populations. Each dot represents an individual tumor. *Statistic:* Unpaired Student's *t*-test.

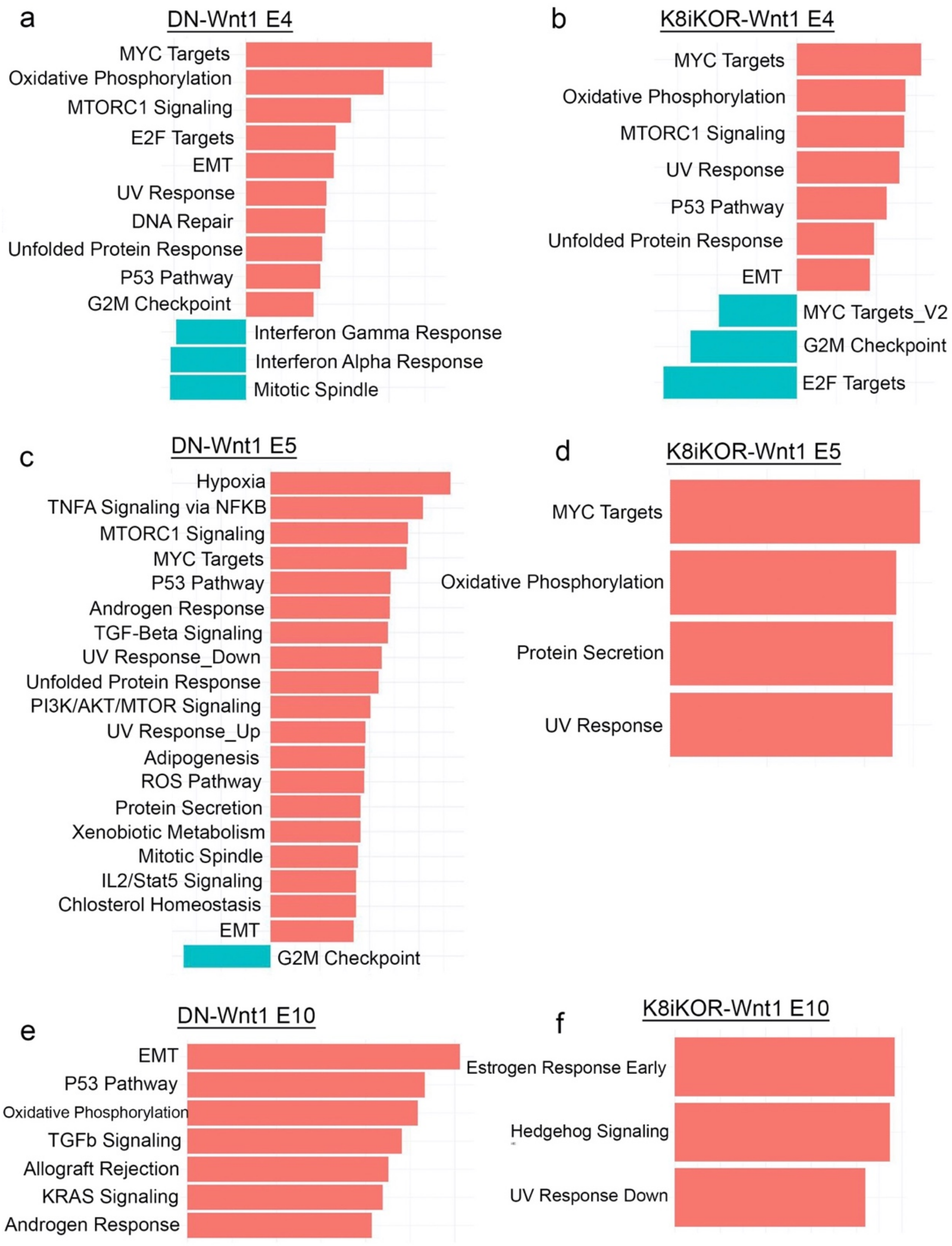


**Supplemental Figure 8: Epithelial-mesenchymal transition (EMT) is enriched in tumor epithelial cells with reduced IGF-1R. a-e.** Fold enrichment bar graphs of the gene set enrichment analysis (GSEA) hallmark gene set in DN-Wnt1 **(a,c,e)** and K8iKOR-Wnt1 **(b,d,f)** primary tumors. Cluster E4 **(a,b)**, E5 **(c,d)**, and E10 **(e,f)** showed enrichment in EMT in tumors with reduced IGF-1R. Red denotes increased fold enrichment, green denotes decreased fold enrichment.

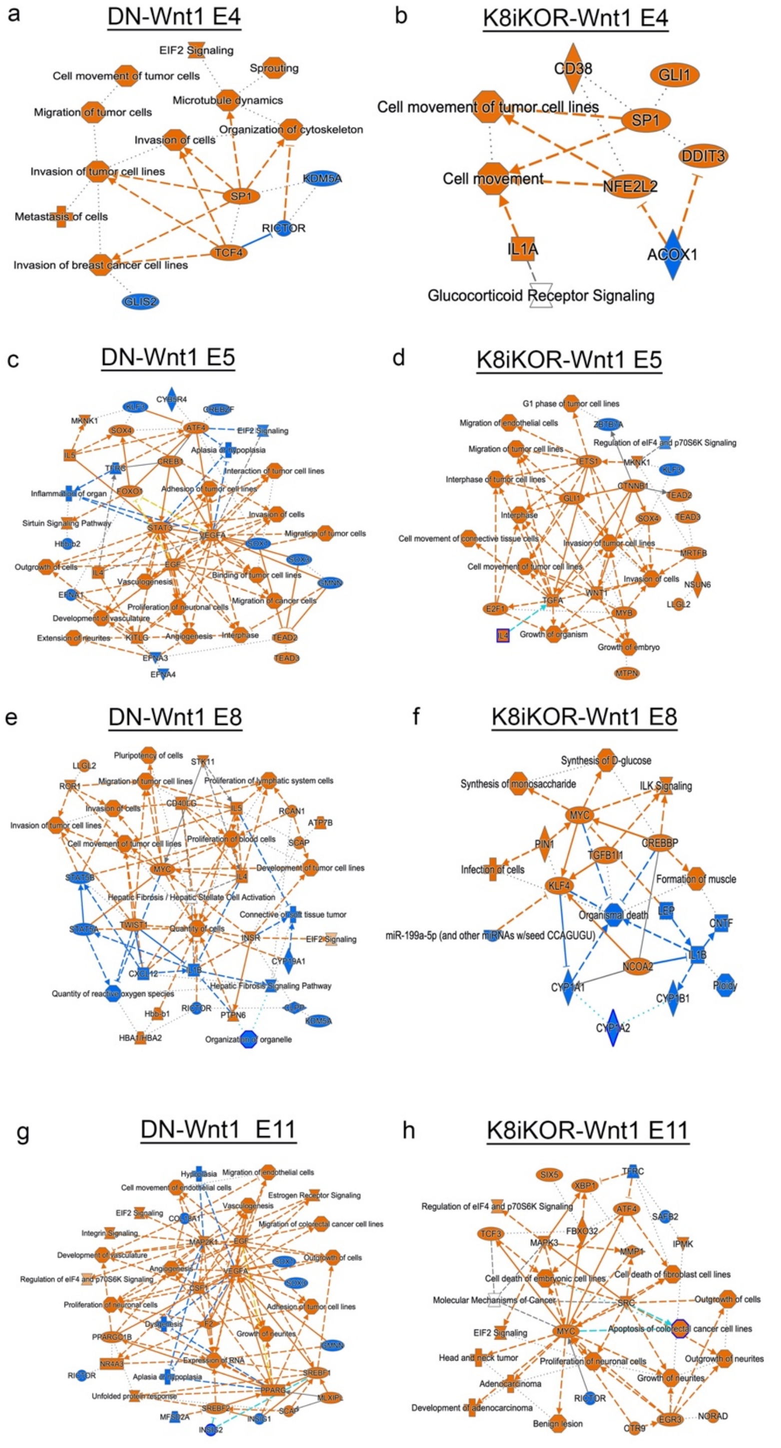


**Supplemental Figure 9: Ingenuity pathway analysis (IPA) of epithelial cell clusters from tumors with reduced IGF-1R compared to Wnt1. a-h.** Graphical summaries from IPA in Cluster E4 **(a,b)**, E5 **(c,d)**, E8 **(e,f)** and E11 **(g,h)** from DN-Wnt1 and K8iKOR-Wnt1 compared to Wnt1 tumors. Blue=downregulated; orange=upregulated.

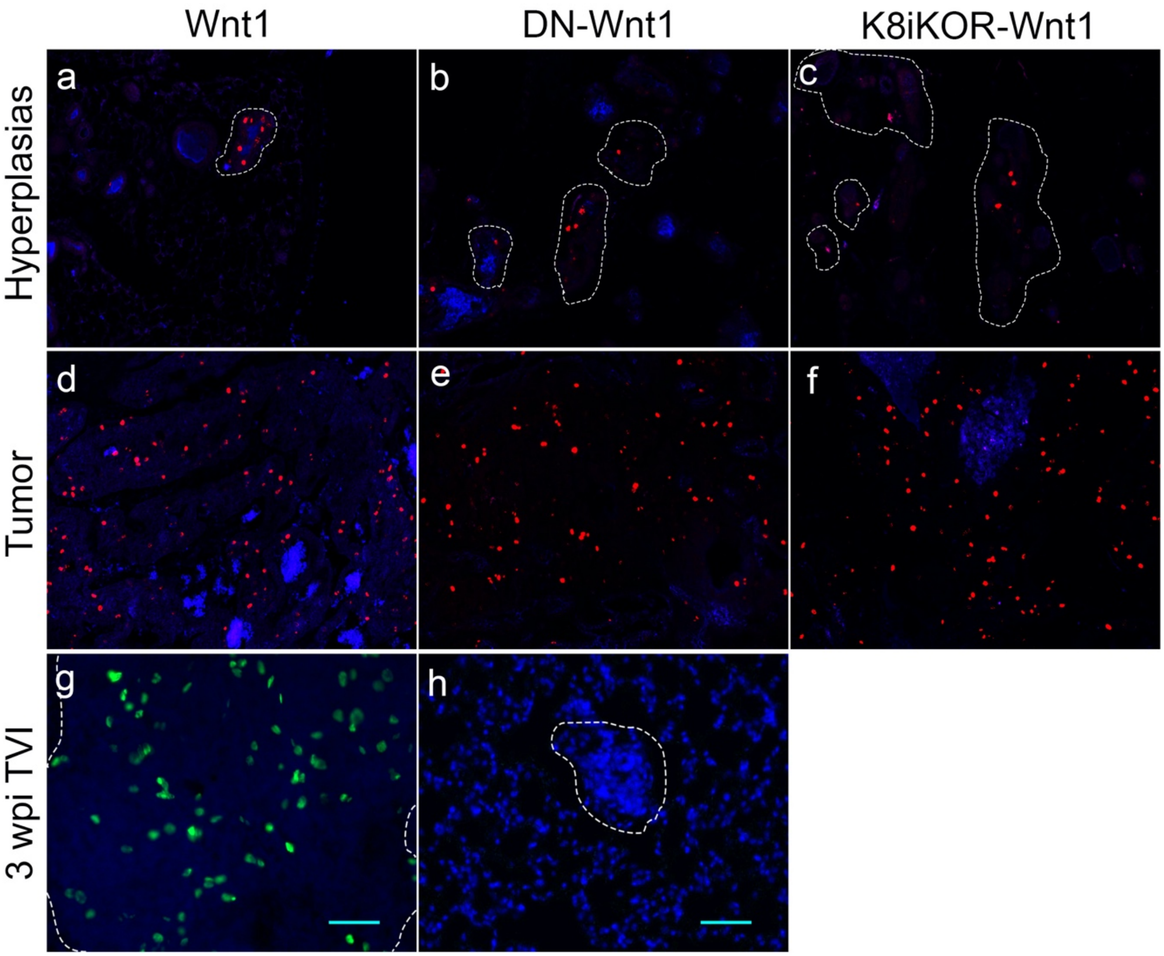


**Supplemental Figure 10: The proliferation rate is unchanged in tumors with reduced IGF-1R compared to Wnt1. a-c.** Representative images of phospho-histone H3 (pHH3) immunofluorescence (IF) in Wnt1 **(a)**, DN-Wnt1 **(b)**, and K8iKOR-Wnt1 hyperplasias. **d-f.** Representative images of pHH3 IF in Wnt1 **(d)**, DN-Wnt1 **(e)**, and K8iKOR-Wnt1 tumors. Red=pHH3 **g-h.** Representative images of pHH3 IF of TVI micrometastases from Wnt1 **(g)** and DN-Wnt1 **(h)** 3 wpi cells. Green=pHH3 Sections are representative of 5 different hyperplasias or tumors and 4 different TVI mets per genotype, scale bar=50 μm.

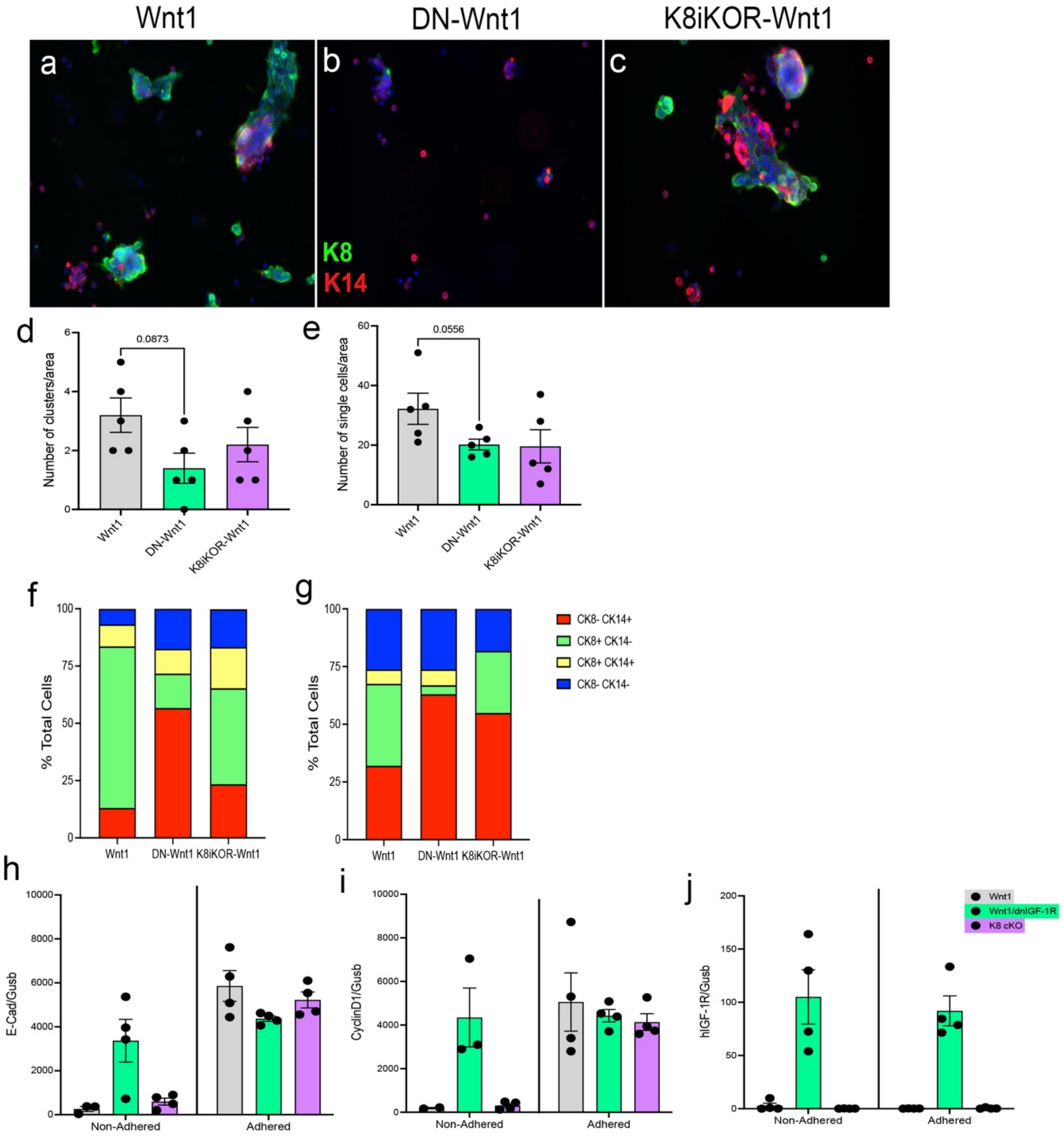


**Supplemental Figure 11: Adhesion is altered in tumors with attenuated IGF-1R *in vitro*. a-c.** Representative images of immunofluorescence for keratin 8 (K8) and keratin 14 (K14) from primary Wnt1 **(a)**, DN-Wnt1 **(b)**, or K8iKOR-Wnt1 **(c)** tumor cells attached to collagen coated plates after 10 hours of incubation. K8=green; K14=red **d-e.** Quantification of clusters **(d)** or single cells **(e)** from primary tumor cells attached to collagen coated plates after 10 hours. *Statistic:* Mann-Whitney U test **f-g.** Quantification of cell types making up the clusters **(f)** or single cells **(g)** from primary tumor cells attached to collagen matrix after 10 hours of incubation. Red=K14$^+$ cells, Green=K8$^+$ cells, Yellow=K8$^+$/K14$^+$ cells, Blue=Keratin negative cells **h-j.** qRT-PCR for E-cadherin **(h)**, cyclin D1 **(i)**, and the *dnIGF-1R* transgene **(j)** in primary tumor cells that did not adhere and that did adhere to collagen after 10 hours.

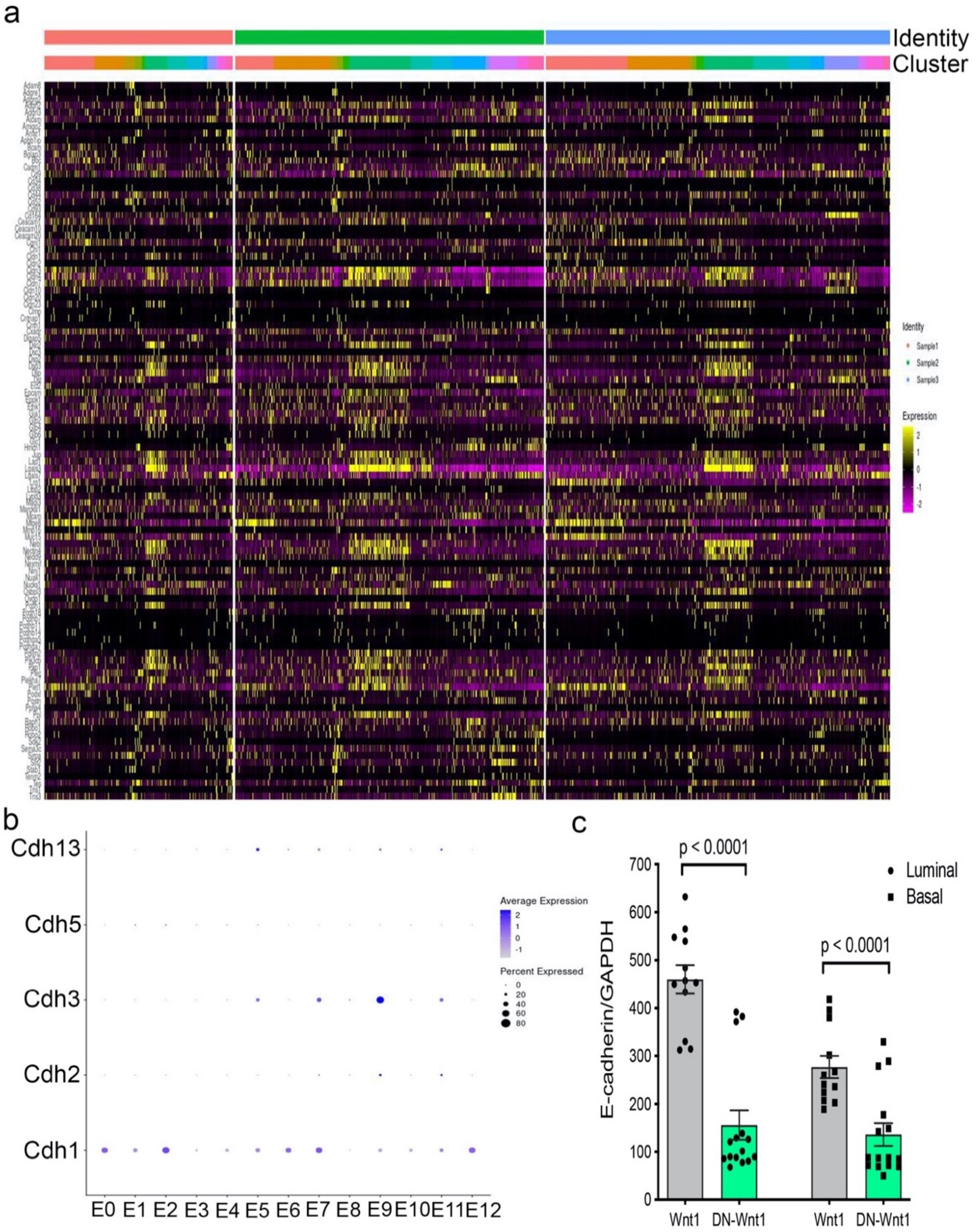


**Supplemental Figure 12: Adhesion gene changes in tumors with reduced IGF-1R. a.** Heat map of adherence genes from single-cell RNA-sequencing. Identity=tumor genotype, cluster=epithelial cell clusters. **b.** Dot plot of cadherins expressed in epithelial cell clusters. **c.** qRT-PCR for E-cadherin from Wnt1 or DN-Wnt1 sorted luminal and basal epithelial tumor cells. *Statistic:* Non-parametric Mann-Whitney U test.

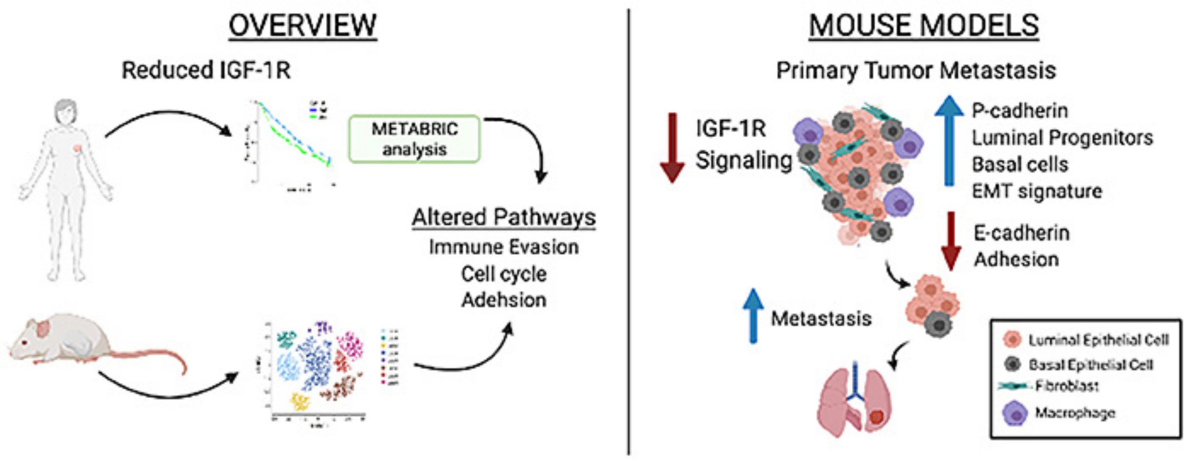


**Supplemental Figure 13. Model for how Reduced IGF-1R in Human Breast Tumors (left) and in Mouse Basal-Like Mammary Tumors results in Enhanced Metastasis.**

| Gene Target | Forward Primer 5' to 3' | Reverse Primer 5' to 3' |
|---|---|---|
| Mouse CyclinD1 | QuantiTect Qiagen Catalog No. 249900 | |
| Mouse E-cadherin | GCTCTCATCATCGCCACAG | GATGGGAGCGTTGTCATTG |
| Mouse GAPDH | GATGCCCCCATGTTTGTGAT | GGTCATGAGCCCTTCCACAAT |
| Mouse Gusb | CAACGCCAAATATGATGCAG | TGCGTCTTATACCAGTTCTCAAAC |
| Mouse IGF-1R | CACAGCTGCAACCACGAG | GGGATATCATCTGCTCCTTCTG |
| Mouse IGF-1R Exon 11 | CGCCTGGAAAACTGCACG | GCAGGGGATACAGTACATGTTT |
| Mouse K14 | GCATCTACTTGCTGGACATCAG | AGGATACCCCAAGCCACTG |
| Human β-Actin | AGCCATGTACGTTGCTATCCA | ACCGGAGTCCATCACGATG |
| Human IGF-1R | GGCACAATTACTGCTCCAAAGAC | CAAGGCCCTTTCTCCCCAC |

**Table 1: List of qRT-PCR primers.**
