## Supplementary material for "Breast tumor Insulin-like growth factor receptor regulates cell adhesion and metastasis: Alignment of mouse single cell and human breast cancer transcriptomics": Obr Supp Methods

### Supplemental Methods

#### K8iKOR-Wnt1 tamoxifen dosage paradigm

The tamoxifen dosage paradigm was determined following a developmental study of the effect of tamoxifen on mammary gland development. Three doses of tamoxifen, 5 mg, 2 mg, 1.5 mg or sesame oil were administered once per day for 3 consecutive days in 4-week-old or 8-week-old FVB mice. Four weeks post-injection, mammary gland development was observed using Carnoy's fixative to clear whole mounted mammary glands. Mammary glands from control samples injected with sesame oil demonstrated no significant changes in secondary or tertiary branching compared to naïve glands, while mammary gland development was stunted with the 5 mg dose of tamoxifen administered at 4 weeks of age. Similar to 4 weeks of age, mammary gland branching was stunted at 8 weeks of age with the 5 mg dose of tamoxifen but not with lower tamoxifen doses. Thus, for all tumor studies, tamoxifen (2 mg for 3 consecutive days) was administered at the end of puberty (8 weeks) to avoid disturbing mammary gland development [20,21] and as confirmed in our studies. Age-matched (8 weeks) females were injected with vehicle sesame oil (control) or tamoxifen for 3 consecutive days to delete the floxed *Igf1r* alleles. Controls for tumor studies included K8-$Cre^{ERT}$ positive females injected with vehicle or K8-$Cre^{ERT}$ negative females injected with tamoxifen. No differences were detected between vehicle and tamoxifen injected controls thus these were combined unless otherwise noted in the methods. Lungs and tumors were harvested when they reached 1.5 $cm^3$. We confirmed deletion of *Igf1r* K8iKOR-Wnt1 by qRT-PCR for *Igf1r* expression (Supp. Fig. 2) and expression of the exon 4 deletion-specific *Igf1r* transcript in tumors and in FAC-sorted luminal epithelial cells.

#### Tumor latency and growth curves

Wnt1 and K8iKOR-Wnt1 female mice were palpated every five days for tumors beginning at nine weeks of age or 1 wpi sesame oil or tamoxifen. Since no differences in latency were observed between vehicle and tamoxifen injected controls, we combined these animals for these studies. Tumor growth was measured by caliper bi-weekly once a tumor was identified, and the mouse was sacrificed when the tumor reached 1.5 $cm^3$.

#### Sorting of mammary tumor epithelial cells

Tumor MECs from either Wnt1 or DN-Wnt1 mice (n=4) were isolated for single cells as described above with minor adjustments for depletion of unnecessary cells. Red blood cells were lysed with a lysis buffer (155 mM $NH_4Cl$, 12 mM $NaHCO_3$, 0.1 mM EDTA) for 5 minutes. Tumor MECs were resuspended at $10^6$ cells/ml in FACS buffer (2% BSA, 2% goat serum in PBS) and immunolabeled with fluorochrome-conjugated cell surface antibodies as described in our previous studies [44]. Single cells were prepared for FACS as previously described [46] and sorted at 70 psi using a 70-um nozzle on the Beckton Dickenson FACS Aria directly into PBS for tail vein injections.

**Flow cytometry analysis of lineage-specific tumor epithelial cells**
Tumor MECs from K8iKOR-Wnt1 mice injected with sesame oil or tamoxifen were isolated for single cells as described above. Since no differences in flow cytometry analysis were observed between vehicle and tamoxifen injected controls, we combined these animals. Tumor MECs were immunolabled with fluorochrome-conjugated cell surface antibodies at $1x10^6$ cells/100ul FACS buffer as described in our previous studies [14,44]. Cells were labelled for viability using a Live/Dead dye (Invitrogen, L34958) and fixed with 1% paraformaldehyde. Single cells were analyzed using the BD LSRFortessa flow cytometer.

**RNAscope analysis of dominant negative IGF-1R expression**
RNAScope Multiplex Fluorescent Assay v2 and a human IGF-1R probe (Advanced Cell Diagnostics, Inc) was used to determine *dnIGF-1R* RNA expression. Tumor tissues were fixed in 4% PFA, paraffin embedded, and sectioned at 7 $\mu$m. Tissue samples were deparaffinized and pretreated with hydrogen peroxide, antigen retrieval, and protease plus reagents. (Mild Reagents Timepoint; RNAScope). Tissue sections were incubated at 40°C (Isotemp Incubator, Fisher Scientific) with either Hs-IGF1R-No-XMm probe (Cat No. 471961), Negative probe (Cat No. 320871), or Positive Probe (Cat No 320881). The probe signal was amplified using Amplification Reagents (RNAScope) and signal was developed using the Multiplex FL v2 HRP-C1, HRP blocker, and Opal 620 fluorophore (Akoya Biosciences, FP1495001KT, 1:3000). Sections were incubated with DAPI (RNAScope) and mounted with ProLong Gold Antifade Mounting medium (Invitrogen). Images were captured on the Keyence BZ-X at 40x and 60x magnification.

**Counting macro and micrometastases in lung sections**
Lung tissue from primary and TVI animals were sectioned at 7 $\mu$m through the entire lung. For coverage of the entire lung, 3 sections were taken and placed on slides and the next 3 sections were disposed through the entirety of the lung tissue or until reaching 72 individual sections. Representative sections (middle section of each 3 sections) were used for H&E staining. Individual macrometastases were counted by eye and micrometastases were counted at 10X magnification with a brightfield microscope (Olympus Provis AX70) from each H&E-stained slide (n=24).

**Single-cell RNA sequencing analysis**
Raw reads were barcode deconvoluted and aligned to the reference genome (mm10) via cellranger (v3.1.0). All subsequent processing was performed using the Seurat package within R (v3.1.5). Low quality cells (cells with percentage of reads of mitochondrial origin >10%, with percentage of reads of ribosomal origin >45%, with <1000 feature counts, with >6000 feature counts) were filtered from the dataset, and read counts were normalized using the scTransform method (https://genomebiology.biomedcentral.com/articles/10.1186/s13059-019-1874-1). Samples were integrated with the Seurat integrate function (https://www.cell.com/cell/fulltext/S0092-8674(19)30559-8), and clustered via UMAP

according to nearest neighbors. Re-clustering was performed as above on subset clusters based on common annotation types.

**WGCNA analysis of METABRIC data for gene module identification**

Patient age and tumor size were coded by ordinal numerical values. Tumor grade was coded as 0 for grade $< 2$ (low) or 1 for grade $> 2$ (high). Cancer subtypes were coded as 0 (TNBC) or 1 (ER+/PR+) resulting in 869 total patients for analysis. IGF-1R expression was represented by IGF-1R Z scores which indicates the number of standard deviations away from the mean of expression in the reference. We selected the top 25% genes with the largest variance of expression values as input for WCGNA, resulting in 12394 genes included in the final analysis.

The association of individual genes with IGF-1R expression was quantified using Gene Significance (GS) as the absolute value of the correlation between the gene and IGF-1R expression. Module membership (MM) was measured as the correlation of the module eigengene (ME) and the gene expression profile. The statistical correlation between modules and trait was denoted and plotted by the correlation and p-values between ME and clinical features. Modules for ingenuity pathway analysis (IPA) (Qiagen) were selected by positive ($> 0.2$) or negative correlation ($< -0.2$) values with significant p-values. IPA was performed as described below.
